# Systematic, Protein Activity-based Characterization of Single Cell State

**DOI:** 10.1101/2021.05.20.445002

**Authors:** Lukas Vlahos, Aleksandar Obradovic, Jeremy Worley, Xiangtian Tan, Andrew Howe, Pasquale Laise, Alec Wang, Charles G. Drake, Andrea Califano

**Affiliations:** Department of Systems Biology, Columbia University Irving Medical Center, New York, NY, USA 10032; Department of Medicine, Columbia University College of Physicians and Surgeons, New York, New York, Columbia Center for Translational Immunology, Columbia University Irving Medical Center, New York, NY, USA 10032; Bristol Myers Squibb, USA 08901; DarwinHealth Inc., New York 10011; Janssen Research and Development, Springhouse, PA, USA; Herbert Irving Comprehensive Cancer Center, Columbia University, New York, NY, US; Department of Biochemistry and Molecular Biophysics, Columbia University, New York, New York; Department of Biomedical Informatics, Columbia University, New York, New York. USA

## Abstract

While single-cell RNA sequencing provides a remarkable window on pathophysiologic tissue biology and heterogeneity, its high gene-dropout rate and low signal-to-noise ratio challenge quantitative analyses and mechanistic understanding. To address this issue, we developed PISCES, a platform for the network-based, single-cell analysis of mammalian tissue. PISCES accurately estimates the mechanistic contribution of regulatory and signaling proteins to cell state implementation and maintenance, based on the expression of their lineage-specific transcriptional targets, thus supporting discovery and visualization of Master Regulators of cell state and cell state transitions. Experimental validation assays, including by assessing concordance with antibody and CITE-Seq-based measurements, show significant improvement in the ability to identify rare subpopulations and to elucidate key lineage markers, compared to gene expression analysis. Systematic analysis of single cell profiles in the Human Protein Atlas (HPA) produced a comprehensive resource for human tissue studies, supporting fine-grain stratification of distinct cell states, molecular determinants, and surface markers.

## Introduction

High-throughput, droplet-based single-cell RNA Sequencing (scRNASeq) has recently emerged as a valuable tool to elucidate the diverse repertoire of cellular subpopulations comprising a broad range of mammalian tissues. Applications of this technology range from study of tissue development^1^ and tumor micro-environment^2^, to the elucidation of tissue heterogeneity^3^ and even of tissue-level response to infectious diseases, such as COVID-19^4,5^. More specifically, scRNASeq data allows identification of representative gene expression signatures for thousands of individual cells dissociated from a tissue sample^6^, thus providing fine-grain characterization of the transcriptional state of individual cell types contributing to the emergence of complex phenotypes, which would be impossible from bulk profiles. This can help elucidate the role of rare populations, for instance, whose gene expression signature would be diluted below detection limits in bulk samples^7^. Moreover, in contrast to flow cytometry or CyTOF, scRNASeq can produce genome-wide single cell profiles, without requiring *a priori* antibody selection and optimization. Not surprisingly, scRNASeq has transformed, virtually overnight, the study of human malignancies, as shown by recent studies in melanoma^8,9^, pancreatic cancer^10^, breast cancer^11^, and renal cell carcinoma^12^.

Despite its many advantages, a key drawback of scRNAseq is that – due to the relatively small number of mRNA molecules per cell and low mRNA capture efficiency - the number of distinct mRNA molecules that can be detected in from each a single cell – as shown by use of unique molecular identifiers (UMI) – is fundamentally limited. This results in scRNASeq profiles that are extremely sparse, with as many as 90% of all genes producing no UMIs in any given cell and the majority of detected genes producing one or two reads. This phenomenon – also known as gene dropout – hinders downstream analyses, challenging quantitative assessment of differential gene expression. As a result, while generating a readout from even 10% of the genes can still provide relatively accurate clustering and classification of broad cell types, many biologically relevant genes – including the established lineage markers of biologically relevant subpopulations – may go undetected. This often results in failure to detect subpopulations associated with more subtle differences such as different fibroblast or macrophage subtypes, which are often defined by the expression of a small set of marker genes^10,12^. Consequently, the ability to identify the mechanistic determinants subpopulation specific cell states is limited, even with when using cutting edge analysis tools such as the Seurat pipeline^13^. Similarly, analysis of genes of interest in single cells is significantly impaired, particularly for transcription factors and signaling molecules that can drive cellular phenotypes even when transcribed at a modest rate.

We have recently developed a paradigm to assess protein activity, defined as the mechanistic contribution of a regulatory or signaling protein to the implementation of a cellular phenotype. Similar to using a highly multiplexed gene reporter assay, we proposed measuring a protein’s activity using a metric based on the enrichment of its transcriptional targets in genes that are differentially expressed in the phenotype of interest, compared to a baseline state. The analysis leverages the ARACNe (Algorithm for Reconstruction of Accurate Cellular Networks)^14,15^ and VIPER algorithms (Virtual Inference of Protein Activity by Enriched Regulon Analysis)^16^. The former is an information theoretic algorithm that has been proven effective in dissecting the tissue-specific transcriptional targets of regulatory and signaling proteins—including ∼6,000 transcription factors, cofactors, signaling proteins, and surface markers—with a relatively modest low false-positive rate (typically in the 20% to 30% range). The latter then computes a Normalized Enrichment Score (NES) representing the enrichment of a protein’s targets in differentially expressed genes. This approach has been used in the literature to identify Master Regulators (MR) proteins, representing mechanistic determinants of cell state, which have been extensively validated^17-20^. VIPER’s reproducibility has led to Dept. of Health approval of OncoTreat^20^ and OncoTarget^21^, two CLIA-compliant clinical tests to predict drug sensitivity of individual tumors.

Because protein activity is assessed based on the expression of *≥* 50 target genes, this methodology is ideally suited for the analysis of single-cell data, where it can provide accurate activity assessment even for proteins whose encoding gene is undetected. The metaVIPER algorithm^12,22^ – specifically developed to extend VIPER to the assessment of protein activity in single cells – has proven effective in stratifying differences between molecularly distinct fibroblast subpopulations^10^, as well as in identifying subpopulations responsible for the presentation of key macroscopic phenotypes, ranging from immune evasion^23^, to relapse following surgery^12^. It has also been instrumental in the identification of MR proteins representing causal, mechanistic determinants of cell state and cell state transitions, such as de-differentiation to a pluripotent stem cell state^24^, transdifferentiation between molecularly distinct, yet co-existing tumor cell states^25^, and beta to alpha cell reprogramming in pancreatic islands^26^.

On an individual basis, these analyses can be quite challenging, due both to the unique complexity and idiosyncrasies of single-cell datasets and to the added complexity of single-cell-based transcriptional network inference. Critically, systematic validation—both technical and biological— across a variety of possible variables, ranging from sequencing depth to number of detected cells, has not previously reported. This provides a rationale for the development of PISCES (Protein Activity Inference for Single Cell Studies), a comprehensive pipeline for the protein activity-based analysis of single-cell scRNA-Seq profiles, made available to the research community as an R package. PISCES leverages the latest implementation of ARACNe (ARACNe3) and leverages NaRnEA (Nonparametric analytical Rank-based Enrichment Analysis)^27^—a more sensitive methodology for gene set enrichment analysis. ARACNe3 implements a more accurate null model based on the principle of maximum entropy (PME) and a much more efficient subsampling procedure, producing higher quality networks from datasets with smaller sample size. Similarly, NaRnEA implements an entirely analytical null model, also based on the PME, which improves protein activity measurement accuracy, while providing separate metrics for statistical significance and effect size. PISCES automates the optimal use of these algorithms for the generation of lineage specific regulatory networks, the identification of molecularly distinct subpopulations, and the identification of MR proteins determining cell state and cell state transitions (Figure 1A).

**Figure 1:**
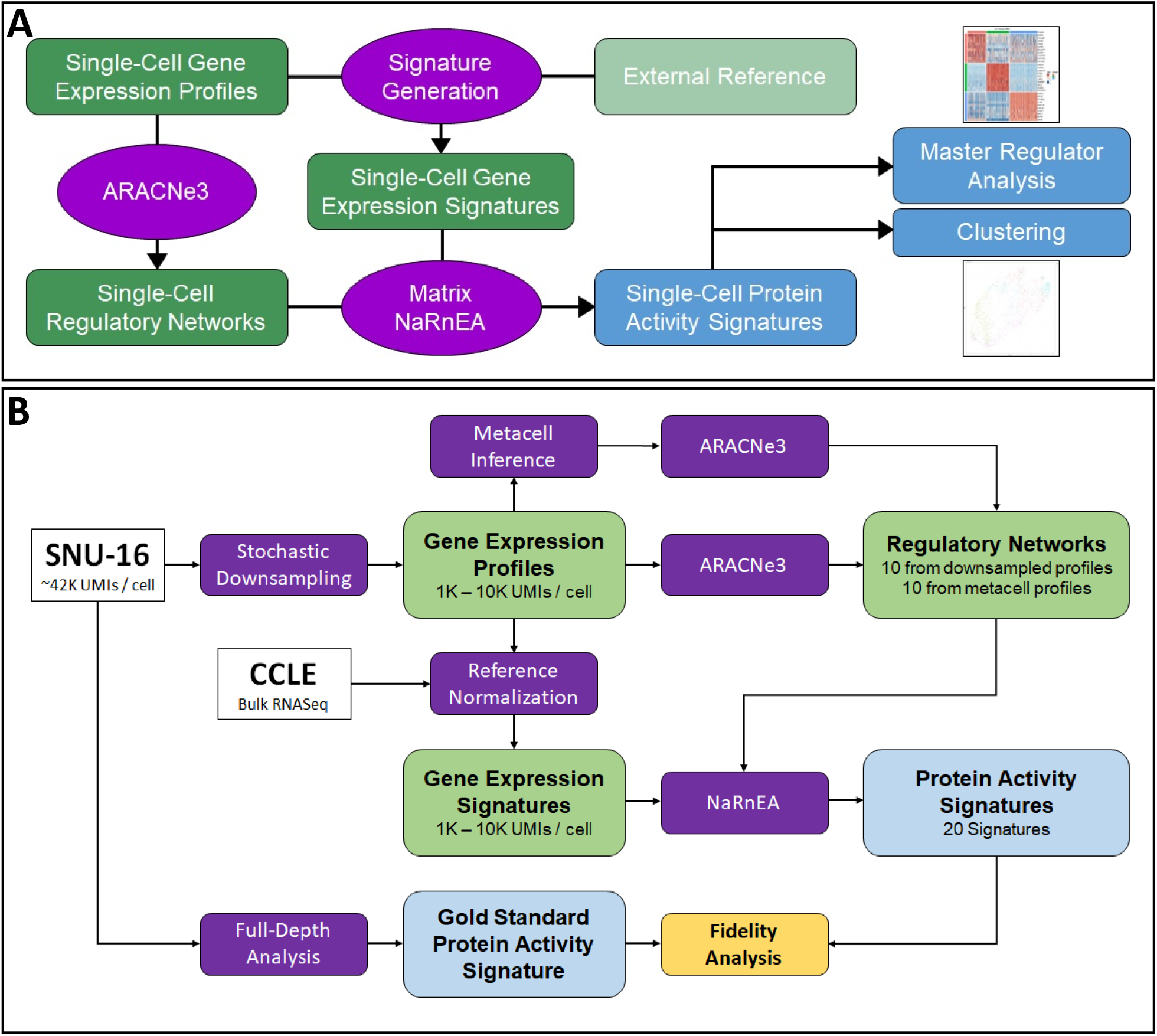
Graphical Representation of Analysis Pipeline and Validation Frameworks

In addition to a comprehensive set of technical benchmarks—for instance to optimize the use of metacell approaches for dataset with low mRNA reads per cell—we provide extensive biological benchmarks for the application of PISCES to the study of complex tissues heterogeneity. These include systematic analyses of Human Protein Atlas (HPA) data^28^—comprising scRNA-Seq profiles from 566,128 cells dissociated from 26 distinct human tissues—as well as 89 tissue-and cell-type specific regulatory networks and MR repertoires, representing a unique resource for the community to study human tissue subpopulations at the single cell level.

Taken together, these analyses show that PISCES significantly outperforms gene expression-based analyses and compares favorably with antibody-based experimental assays, while supporting proteome-wide quantitative analyses. Thus, PISCES provides a valuable complement to existing methodologies for the analysis of single-cell datasets, which greatly improves the granularity of cell subpopulation detection, supports the characterization of biologically relevant subpopulations that may be missed by gene expression-based approaches due to gene dropout, and allows for the accurate assessment of Master Regulators of single-cell states.

## Results

### PISCES Overview

PISCES takes a single-cell Unique Molecular Identifier (UMI) count matrix as input, with rows representing genes and columns representing different cells. Initial quality control (QC) is performed based on cell depth, number of detected genes, and mitochondrial read fraction in each cell; the parameters for these analyses should be adjusted based on the specific tissue being analyzed and the library preparation methodology. Gene expression is then normalized by converting counts to Log2(CPM + 1), where CPM indicates counts per million.

Following normalization, an initial gene expression-based clustering is performed, using the preferred unsupervised clustering methodology (e.g., via Seurat analysis) or through the use of an external annotation tool such as SingleR^29^. The goal of this step is to organize cells into broad, lineage-specific subset for network generation. This is important because we have shown that, due to consistent or divergent epigenetic cell state, regulatory networks are highly conserved within closely-related cell lineages^30^ and yet quite distinct when compared across lineages^31^. As a result, such a step is not necessary when analyzing homogeneous datasets, comprising cells with a largely conserved regulatory architecture (*e*.*g*., a FACS-purified T cell population).

Once lineage-specific clusters have been generated, ARACNe3 is used to generate networks for all adequately sized clusters, considering both sample size and sequencing depth (see technical validation). For datasets with < 10,000 UMIs/cell, metacells are generated by aggregating unnormalized counts from multiple cells using a k-nearest neighbors approach. While the default is k = 5, this parameter should be adjusted for optimal results to generate metacells with *≥* 10,000 UMIs/cell. To reduce metacell overlap, a subset (typically n = 250) of all possible metacells should be used, which is then Log2(CPM + 1) normalized and used as input to ARACNe3. Three different regulatory protein sets are used, including (a) transcription factors (TFs) and co-factors (co-TFs), (b) signaling proteins, and (c) surface proteins. These sets can be modified as needed or excluded from the analysis based on the specific biological question. PISCES provides such predefined sets for both human and mouse studies and additional organism-specific sets can be added as needed.

Differential gene expression signatures are then generated from the normalized gene expression counts, by comparing the scRNA-seq profile of each cell either to the centroid of the entire populations (as a default) or to an external, user-provided reference (see STAR methods). These signatures and all cluster-specific networks are then used as an input to a refined version of the metaVIPER algorithm (metaVIPER2)—which leverages NaRNeA as an improved core gene set enrichment analysis algorithm—to measure protein activity (see STAR methods). MetaVIPER2 further improves over the original metaVIPER implementation by using a probabilistic framework to weight the contribution of different networks to protein activity measurement, based on their statistical match to the specific cell being analyzed, on a protein-by-protein basis (see methods). NaRnEA produces both a Normalized Enrichment Score (NES) as a measure of statistical significance and a Proportional Enrichment Score (PES) as a measurement of effect size. The latter is not affected by the number of targets of a protein and is thus used as an unbiased assessment of protein activity for subsequent analyses.

PISCES provides a variety of clustering methodologies for both discrete (silhouette-score optimized Louvain clustering) and continuous (multi-way K-means) clustering. Additionally, it includes several methods for the identification of MR proteins, whose differential activity best separates distinct subpopulations. The standard option uses a Kruskal-Wallis test to identify proteins that best differentiate cluster identity, but other solutions—e.g., based on Cohen’s Kappa, group-specific Wilcoxon Rank Sum test, or cluster specific integration—are also included (see STAR Methods). PISCES include appropriate visualization methods for the generation of cluster heatmaps and violin plots, see schematic workflow representation (**Figure 1A**).

### Benchmarking Overview

For benchmarking purposes, including optimal parameter selection and default options, we leveraged publicly available PBMC CITE-Seq data. Overall, we tested three normalization methods, four signature generation methods, three cluster methodologies for network generation, and two methods for protein activity assessment. Optimal parameter selection was assessed based on the concordance between antibody-based abundance and metaVIPER-assessed protein activity. See STAR Methods and Suppl. Table S2 for a comprehensive description of parameter space exploration.

It should be noted that, due to intrinsic differences across multiple single-cell datasets, no parameter choice is universally optimal. In particular, we recommend that the clusters utilized for network generation be carefully vetted and that a final selection be made based on the underlying dataset biology. For this purpose, PISCES allows parameters to be set by the user and its modular implementation allows substituting any of the provided methodologies, such as the preferred clustering algorithm, with user-defined ones.

### PISCES provides highly reproducible protein activity assessment from low-depth single cell profiles

To determine whether PISCES-measured protein activity provided more robust signal for low depth data, we performed technical validation using the SNU-16 cell line, a relatively homogenous stomach adenocarcinoma model that is transcriptionally complex and produces high UMI counts (i.e., 41,915 UMI/cell across 6,157 single cells). ScRNA-seq profiles were then stochastically subsampled reads in each cell to depths between 1,000 and 10,000 UMIs at 1,000 UMI intervals (see Methods). Read-subsampled data were Log2(CPM + 1) normalized and gene expression signatures were generated by comparing each profile to the centroid of the Cancer Cell Line Encyclopedia (CCLE) from the Broad^32^, as a reference control, read-subsampled via the same procedure to match the test sample’s depth (see Methods). Additionally, to assess metacell-based performance, we generated ARACNe3 networks from each read-subsampled dataset with and without metacell assembly (see Methods).

As a “gold standard” reference for reproducibility, we assessed protein activity using the full-depth scRNA-seq profiles. Specifically, to ensure optimal network generation quality, we analyzed the full-depth gene expression profiles with ARACNe3, at FDR = 0.05. We then used metaVIPER2 to generate protein activity profiles from the full-depth gene expression profile of each single cell.

To benchmark PISCES, we performed three separate analyses. First, we measured protein activity from read-subsampled gene expression signatures, using the full-depth network. Second, we measured protein activity using both read-subsampled gene expression signatures and networks generated from read-subsampled profiles, without metacells. Finally, we repeated the latter analysis using metacells.

To establish robustness of protein activity assessment as a function of scRNA-seq profiling depth, we used NaRNeA to assess the enrichment of the 50 most differentially active proteins from the full-depth analysis (FDR < 0.05), weighted by their rank (association weight) and differential activity sign (association mode), in proteins differentially active based on each of the subsampled analyses (**Fig. 2a**). The same metric was used to measure robustness of gene expression assessment, based on the enrichment of the 50 most differentially expressed genes.

**Figure 2:**
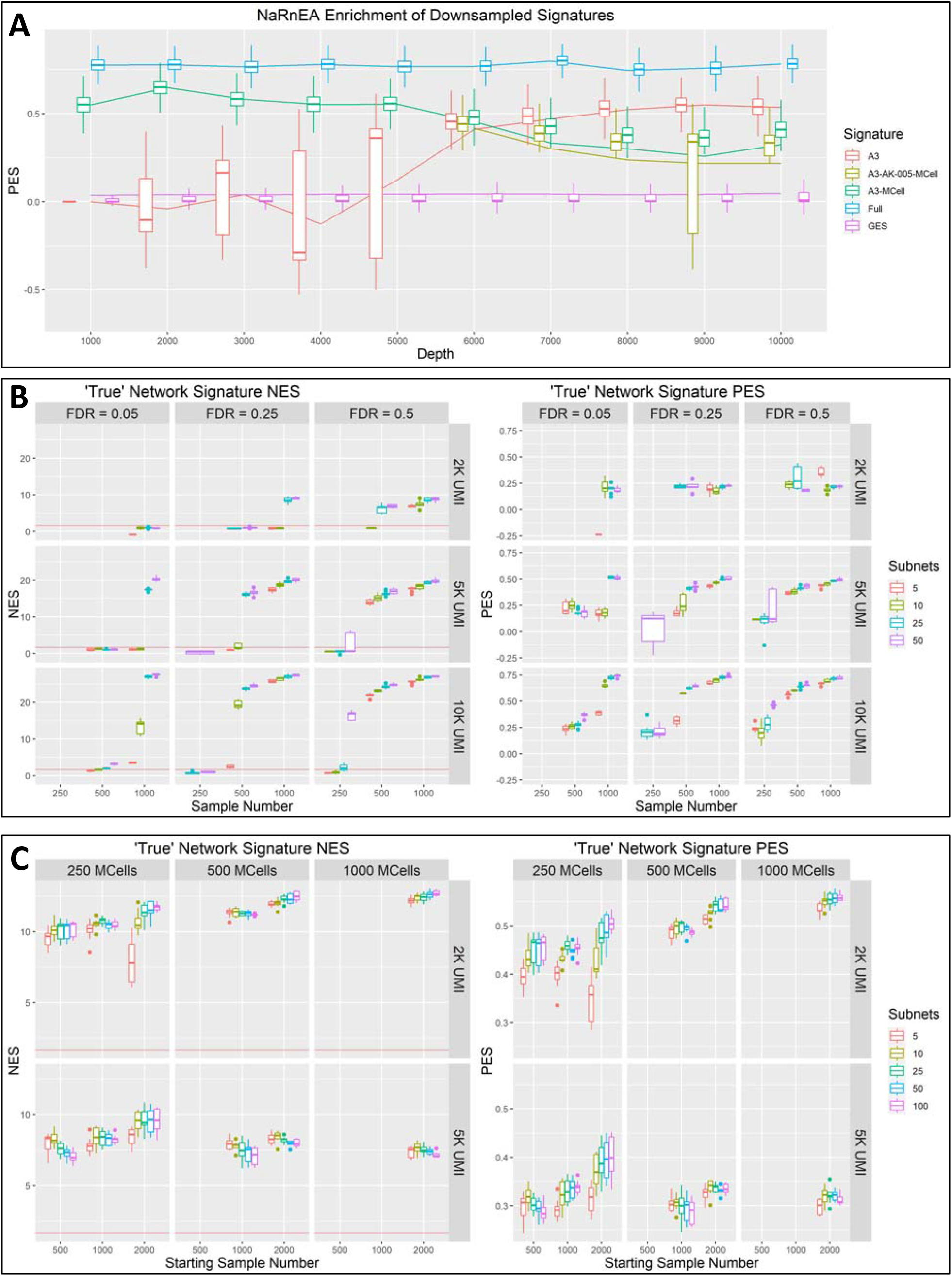
Technical Benchmarking Shows Increased Recover of Original Data Structure from Read-Subsampled Matrices by PISCES vs Gene Expression **2A)** Sample-specific Proportional Enrichment Score (PES) distributions for the enrichment of full-depth regulons in read-subsampled data. Read-subsampled protein activity signatures are generated using a read-subsampled gene expression signature and either a non-metacell network (red), metacell network (green), or the full-depth network (blue). Enrichment of the full-depth gene expression signature regulons in read-subsampled gene expression signatures are shown in purple. **2B)** Group-specific PES and Normalized (NES) distributions for the enrichment of full-depth signatures in protein activity signatures generated from parameter-permuted networks. Parameters tested are the null-model FDR (columns), starting sample depth (rows), number of samples (x-axis), and the number of subnetworks (colors). The red line corresponds to an FDR corrected p-value of 0.05. **2C)** Group-specific PES and Normalized (NES) distributions for the enrichment of full-depth signatures in protein activity signatures generated from parameter-permuted metacell networks. Parameters tested are the number of metacells (columns), starting sample depth (rows), number of starting samples prior to metacell generation (x-axis), and the number of subnetworks (colors). The red line corresponds to an FDR corrected p-value of 0.05.

When using the full-depth network, differential protein activity was dramatically more reproducible compared to differential gene expression. This shows that high-quality networks can help reproducibly assess protein activity, even from samples with as few as 1,000 UMI/cell, because protein activity is assessed from the expression of a large number of high-confidence regulatory targets. Indeed, a high-quality network allows measuring the activity even of proteins encoded by genes that, due to gene drop out, have no mRNA reads.

Among the genes encoding for proteins analyzed by metaVIPER2, > 33% lacked adequate reads across the entire dataset (Table S4). In sharp contrast, metaVIPER2 analysis, using the full-depth network, could measure the activity of all selected proteins, independent of read subsampling up to 1,000 UMI/cell. Moreover, metaVIPER2 analysis could assess statistically significant differential activity for thousands of proteins, while gene expression could only identify a few hundred genes as significantly differentially expressed in at least 5% of cells across all subsampled depths (Table S5) (see Methods).

While high-quality networks will become increasingly available, including as a result of these studies, they may still be unavailable for some cell lineages. In this case, networks will need to be generated from potentially low-depth scRNA-seq data. Our analyses show that, for sequencing depths between 5,000 and 10,000 UMIs, reproducibility of protein activity signatures from read-subsampled data (without the use of metacells) significantly outperform that of gene expression. Reproducibility of protein activity at lowed sampling depths, using metacell-based networks, also significantly outperformed gene expression reproducibility. A full report of log p-values for the comparison of PES distributions between each protein activity signature’s ground truth enrichment and that for the corresponding gene expression enrichment can be found in Table 1.

**Table 1:**
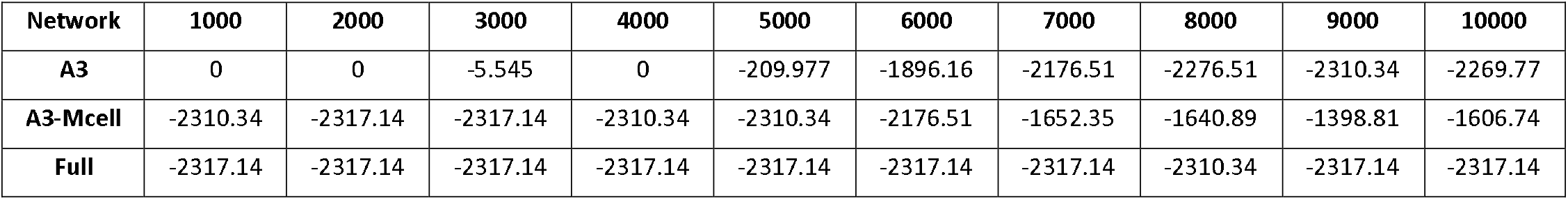
LogP values for concordance of subsampled protein activity w/ full depth

### Selecting optimal ARACNe3 analysis parameters

We leveraged the same subsampling strategy to explore optimal parameter selection for ARACNe3 network generation. Specifically, we assessed optimal selection of two parameters, i.e., the number of bootstraps (n = 5, 10, 25, and 50 networks) and mutual information statistical significance threshold (FDR = 0.05, 0.25, and 0.5), as a function of two variables, including both sequencing depth (n = 2,000, 5,000, and 10,000 UMI/cell) and sample size (n = 250, 500, and 1000 samples). For each of the 108 possible choices, we generated ARACNe3 networks (without metacells) and read-subsampled gene expression signatures by compared each sample to the depth-matched, read-subsampled CCLE centroid. These were then used to assess both protein activity and gene expression signature reproducibility to the corresponding profiles generated from the reference dataset (see Methods). Statistically significant reproducibility was assessed at p ≤ 0.05, as assessed from the enrichment analysis’s NES of the top 50 proteins/genes; overall reproducibility effect size was assessed based on PES (Figure 2B).

These analyses show that ARACNe3 network generation is highly robust to parameter selection. For instance, even as few as 2,000 UMI/cell, high reproducibility protein activity profiles could be generated with the appropriate combination of FDR threshold, bootstrap number, and sample size. Interestingly, as previously reported, using permissive FDR values does not produce noticeable decrease of protein activity reproducibility^27^ because interactions that are not statistically significant are eliminated by maximum entropy pruning step in ARACNe3.

### Metacells further improve reproducibility from low-depth profiles

To assess whether metacells may improve reproducibility of low-depth analyses and to further explore the parameter space, we repeated the previous section’s analysis for the 2,000 and 5,000 UMI/cell read-subsampled datasets, using k = 5 metacells and FDR = 0.25. We measured the effects of using a different numbers of metacells (n = 250, 500, and 1,000) and different dataset sizes (n = 500, 1,000, and 2,000 single cells). Reproducibility was assessed as previously described.

The analysis shows that, even when a limited number of low UMI-depth cells are available, use of metacells can produce networks that generate high-reproducibility protein activity signatures (**Figure 2C**). Moreover, the subsampling strategy used for metacell generation effectively removed the potential adverse effects that may result from the use of partially overlapping metacells.

Critically, the analysis provides an effective guideline to determine when metacells should or should not be used to improve network generation from low-depth datasets. Specifically, metacells significantly improved reproducibility for datasets with < 5,000 UMI/cell. For these, networks should be generated using metacells with k = 3 to 5 and a permissive FDR = 0.25, with more single cells producing better quality networks, with smaller gains for datasets with 5,000 to 7,000 UMI/cell. In sharp contrast, non-metacell networks provide better reproducibility for datasets with *≥* 7,000 UMI/cell.

### Protein activity outperforms gene expression based on antibody measurements

While the technical benchmarks provide effective evidence on reproducibility, they cannot assess whether PISCES-based activity assessment is consistent with experimental assays. To address this challenge, we assessed concordance of PISCES-measured protein activity with CITE-Seq based protein abundance, using a publicly available dataset of peripheral, blood-derived mononuclear cells (PBMCs)^33^. See Suppl. Table 1 and Suppl. Figure S1a for relevant QC metrics. This dataset was analyzed with the standard PISCES protocol, using Log2(CPM + 1) normalization, internal scaling, and SingleR-derived lineage networks with k = 5 metacells, and protein activity measured by MetaNaRnEA.

To assess the statistical significance of the agreement between antibody-derived tags (ADT) measured by CITE-Seq and either protein activity or gene expression, we used the Dunn test, as implemented in ‘cocor’ R package^34^ (see Methods). The analysis shows highly significant improvement (p = 6.5×10^−968^) in Spearman’s correlation between ADT and protein activity (ρ = 0.41) compared to gene expression (ρ = 0.14) (Figure 3A). Recognizing that protein activity and protein abundance are not identical, it is reasonable to expect that the true assessment of protein activity should be even better than assessed by this test.

**Figure 3:**
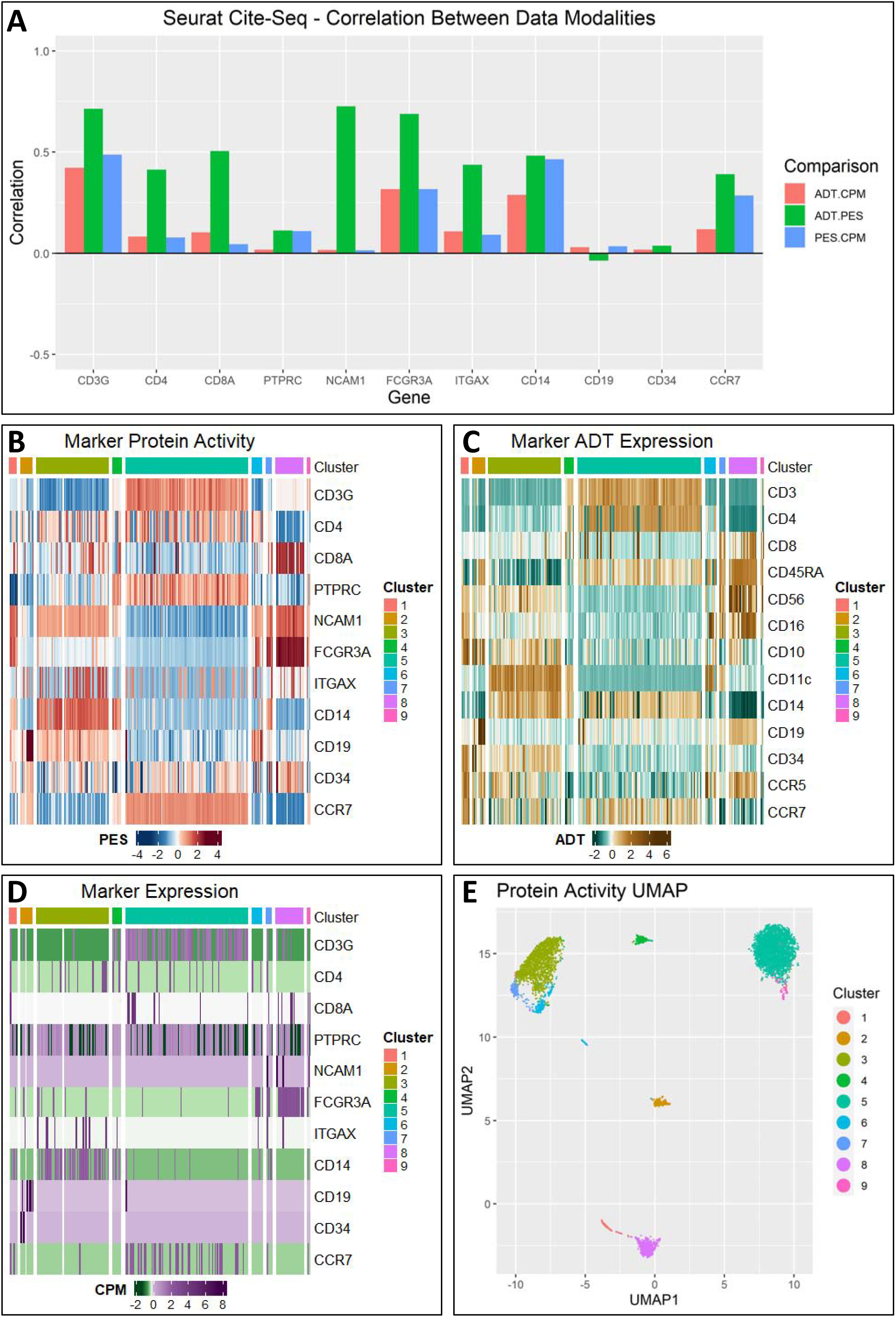
Biological Benchmarking Shows Dramatically Increased Concordance with CITE-Seq Antibody Profiling by PISCES vs Gene Expression **3A)** Spearman correlation between antibody abundance (ADT) and Log(CPM + 1) normalized gene expression (CPM) (red), ADT and proportional enrichment score (PES) derived from PISCES (green), and CPM and PES (blue) in the Seurat PBMC CITE-Seq dataset. PISCES-measured protein activity significantly outperforms gene expression for the majority of proteins. **3B-D)** Protein activity-derived clustering displaying marker activity, antibody abundance, and marker expression respectively. Marker segregation is significantly clearer and more concordant with antibody abundance levels at the activity level relative to expression. **3E)** Protein activity-derived UMAP, colored by clusters.

We also used this dataset to perform the parameter exploration described in previous sections, see Table 2 for the best parameter selection based on the Dunn metric and Suppl. Table S2 for the complete set of results.

**Table 2:** Top PISCES Paramaters by improved concordance w/ antibody abundance relative to gene expression

| Signature | Network | Method | Zscore | logp | ADT.CPM | ADT.PES | num.genes |
| --- | --- | --- | --- | --- | --- | --- | --- |
| cpm-scale | sr-lineage | meta-narnea | 66.66205731 | -2227.033742 | 0.140183999 | 0.406365592 | 11 |
| pflpf-scale | sr-lineage | meta-narnea | 65.32314159 | -2138.654932 | 0.145498756 | 0.405773932 | 11 |
| pflpf-scale | sr-lineage | net-match-narnea | 52.48451224 | -1382.191832 | 0.141554294 | 0.344657347 | 11 |
| cpm-scale | sr-lineage | net-match-narnea | 52.24318067 | -1369.550177 | 0.142665497 | 0.342242829 | 11 |
| cpm-ecdf | sr-lineage | meta-narnea | 46.96238124 | -1107.501364 | 0.136664084 | 0.320177143 | 11 |
| sct-full | sr-lineage | meta-narnea | 46.60165866 | -1090.61833 | 0.143478972 | 0.333362369 | 11 |
| sct | sr-lineage | meta-narnea | 46.00432983 | -1062.947327 | 0.144591397 | 0.332587728 | 11 |
| pflpf-ecdf | sr-lineage | meta-narnea | 45.63620318 | -1046.07164 | 0.137136496 | 0.317123559 | 11 |
| sct | sr-lineage | net-match-narnea | 39.22133767 | -773.7454727 | 0.143991578 | 0.30352525 | 11 |
| cpm-pseudo-ecdf | sr-lineage | net-match-narnea | 38.68524913 | -752.849314 | 0.138907128 | 0.306889995 | 11 |

To ensure that these results were not idiosyncratic, we repeated the analysis using an independent PBMC CITE-Seq dataset (see Methods). This data had significantly higher depth, allowing us to assess the performance of PISCES when using smaller, k = 2 metacells. Consistent with the first analysis, we observed highly significant increase (p = 1.5×10^−91^) in correlation between ADT and protein activity (ρ = 0.351) compared to gene expression (ρ = 0.273) (**Figure S2A**).

Finally, to further assess improvement in low-depth datasets, a strength of the methodology, we repeated the analysis after stochastically subsampling the second CITE-Seq data to 2,500 UMI/cell and analyzing the read-subsampled profiles using k = 5 metacells, as suggested by the parameter exploration. As expected, the increase in correlation between ADT and protein activity was dramatically increased compared to gene expression, (ρ = 0.307 vs. 0.185, p = 4.5×10^−243^) (**Figure S3A**). These findings further confirm that PISCES may provide an effective tool for the study of single cell datasets.

### PISCES improves subpopulation stratification of complex tissues

To further establish the biological value of PISCES, we compared activity-and expression-based clusters identified from the Seurat PBMC CITE-Seq dataset. Cluster quality was assessed by two complementary metrics, including (a) silhouette score, based on a distance metric assessed from ADT levels and (b) and a Kruskal-Wallis test of the clustering’s ability to segregate ADT markers (see STAR methods). Both provide effective metrics to assess whether activity-based clusters effectively co-segregate with ADT marker-based clusters, compared to gene expression analysis.

Cluster analysis for both protein activity and gene expression was performed using the silhouette-optimizing Louvain algorithm, which is part of the standard PISCES analytical pipeline. We tested four distance metrics; Euclidean, spearman correlation, PCA-Euclidean, and PCA-correlation (spearman), all measured from the protein activity space. Metrics for all parameter permutations are presented in Supplementary Table 3.

This analysis finds highly statistically significant segregation of ADT marker genes by Kruskal-Wallis test (p-value < 2.2e-16) for all parameter permutations, independent of distance metric selection. Moreover, activity-based clusters generated using Euclidean and Correlation distance metrics both show silhouette scores > 0.25, a generally accepted threshold for meaningful clustering^35^ (Suppl. Table 3). Visualization of activity, expression, and antibody-based marker gene measurement confirmed these results. Indeed, while cell lineage stratification based on the activity of established markers is rather obvious, gene expression produced a much noisier result due to high gene dropout (**Figure 3B-D**). Interestingly, while most gene expression-based analyses benefit greatly from feature reduction before distance calculation—*e*.*g*., based on variance analysis—this step is virtually unnecessary for protein activity-based analyses.

Unsupervised, activity-based cluster analysis identified 9 subpopulations, with distinct activity of canonical lineage markers (**Figure 3B**). Overall, protein activity profiles were highly consistent with measured ADT protein abundance, with activity successfully measured for 12 out of 14 marker proteins (**Figure 3C**), In contrast gene expression is too noisy to provide useful interpretation for the majority of markers, with only five genes having interpretable patterns (**Figure 3D**). CD4 T-cells (cluster 5 and cluster 9)—identified by high activity and abundance of CD3 (CD3G), CD4, and CCR7 marker proteins—represent the most abundant subpopulation. CD8 T-cells (cluster 8) are clearly identified by high activity of the protein encoded by the CD8B gene and ADT-measured abundance of CD8 protein. An otherwise phenotypically similar population negative for CD8 (cluster 1) is likely comprised of Natural Killer cells. Cluster 2 is significantly positive for CD19, a canonical B-cell marker, while clusters 3 and 7 are both strongly enriched for protein activity of the canonical myeloid marker CD14, although only cluster 7 is also positive for CD16 (FCGR3A). Thus, these clusters match the canonical classification of CD14+CD16-classical monocytes and CD14+CD16-non-classical monocytes^36^. Of note, cluster 4 and cluster 6 show high activity of proteins associated with multiple cell identities: cluster 4 is positive for both the CD4 T-cell markers (CD3/CD4) and the myeloid marker CD14, while cluster 6 is positive for CD3/CD4 and the B-cell marker CD19. As such, these clusters are likely to represent doublets comprising CD4 T-cells and either a myeloid or a B-cell, respectively.

When repeated on both the original and subsampled CITE-Seq from the Sims lab, the analysis revealed similar cluster structure, showing clear segregation of CD4 T-cells (clusters 1 and 6), CD8 T-cells (clusters 2 and 5), and myeloid cells (clusters 3 and 4) (**Suppl. Figure s2B-E, s3B-E**).

### Analysis of the Human Protein Atlas

Validation studies on CITE-Seq data establish that protein activity-based clustering outperforms the same analysis using gene expression. This increased ability to stratify single cells into biologically meaningful subpopulation is most effectively demonstrated by performing systematic analyses of a large collection of single cell profiles from multiple human tissues. For this purpose, we performed comprehensive analysis of scRNAseq profiles from the Human Protein Atlas (HPA), for a total of 566,128 cells dissociated from 26 distinct human tissues. The analysis required generation of 88 lineage/tissue-distinct networks. These represent a unique community-available resource to perform single cell analyses to elucidate human tissue heterogeneity and mechanisms from any novel datasets. In addition, for each tissue we performed tissue-level activity-based clustering—identifying 224 tissue-specific clusters—as well as Master Regulator analysis, identifying proteins representing candidate mechanistic determinants of subpopulation-specific transcriptional states.

To demonstrate PISCES’s ability to recapitulate the expected activity patterns of canonical subpopulation markers, we selected two tissues for more in depth assessment. First, we studied subpopulations identified by unsupervised activity-based cluster analysis of the HPA PBMCC dataset. As expected, even though antibody-based data was not available, the analysis reproduced the biologically relevant clusters that had emerged from CITE-Seq datasets (**Figure 4A-B**), including clear identification of CD4+ (clusters 1, 4 and 6) and CD8+ T-cells (clusters 3 and 7), as well as CD19+ B-cells (cluster 5), and CD16(FCGR3A)+ myeloid cells (clusters 2 and 3). Then, to further demonstrate the ability to identify established markers in unsupervised fashion, we studied subpopulations and associated MR proteins identified in the HPA Kidney dataset, showing robust separation of known subpopulations in the renal epithelium (Figure 4C).

**Figure 4:**
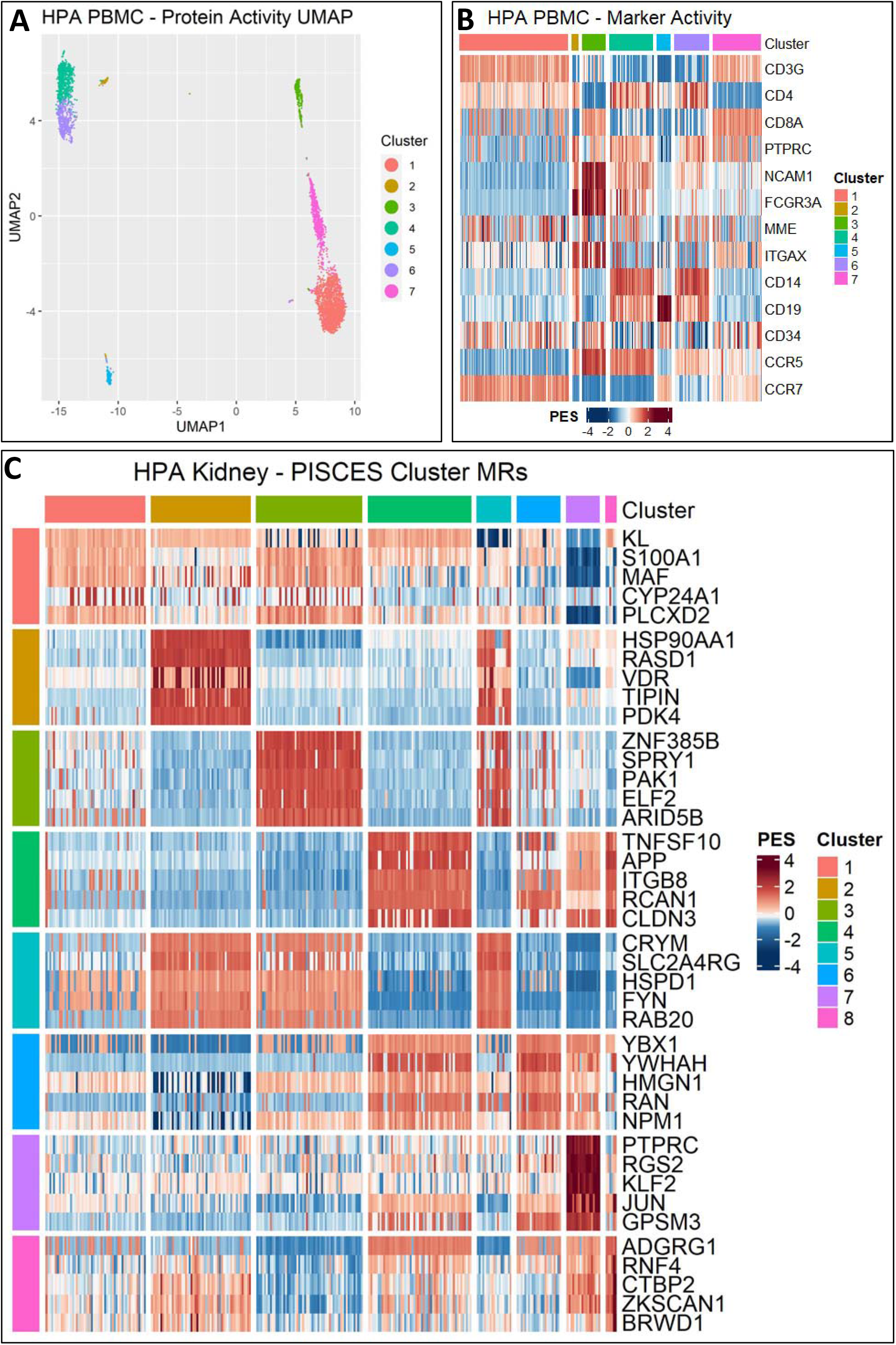
PISCES Analysis of the Human Protein Atlas with exemplar results from PBMC and Kidney **4A-B)** Exemplar results from the HPA PBMC data, with clusters generated from protein activity represented in UMAP space and the activity of classical PBMC markers. This unsupervised analysis identifies canonically expected immune cell subtype, providing a positive control for the robustness of protein activity-based clustering analysis. **4C)** Exemplar results from the HPA Kidney data, with the activity of cluster-specific master regulators as determined by the Kruskal-Wallis MR procedure. Here again we see many classically observed markers of Kidney cell subtypes, including a resident immune population. Further work would explore the novel MRs identified for each of these clusters.

Among non-epithelial cells, the analysis identified a well-separated population of tissue-resident immune cells (cluster 7), marked by high activity of PTPRC (CD45) and CD5, as well as other canonical immune-cell-specific proteins; further sub-clustering, to identify specific subpopulations of immune cells, was limited by cell count. Within the epithelial compartment, three clusters of proximal tubular epithelial cells were identified (cluster 1, 4, 8), where cluster 1 is defined by high activity of Klotho (KL), a known cofactor of FGF23 highly active in proximal renal epithelium^37^, and clusters 4 and 8 also exhibit high activity of TNFSF10, a key regulator of epithelial-to-mesenchymal transition in proximal epithelium^38^. Clusters 2, 3 and 5 are phenotypically similar and likely represent distal tubular epithelial cells, each exhibiting high activity of CRYM, known to be highly represented in the renal medulla^39^. Finally, cluster 6 is shares protein activity patterns with the other epithelial populations, especially clusters 4 and 8, and is further defined by high differential activity of HMGN1 and YBX1, known regulators of renal tubular inflammatory injury via the NfKB signaling pathway^40^. Overall, robust, biologically consistent clustering is achieved by PISCES on renal and blood tissues, suggesting that high-quality stratification is also achieved for the other tissues, thus making an extensive resource available to the community.

Finally, we extended the analysis of the HPA tissues in a hierarchical manner, performing subcluster analysis of each lineage identified by the initial PISCES analysis. Initial lineage classifications were determined by HPA’s internal labeling or by SingleR when HPA’s labels were anatomical rather than cell-type specific. This analysis generated 492 subclusters across all tissues, for each of which we also performed unsupervised Master Regulator analysis. As an example, we show the result of subclulstering the lymphoid cell cluster identified by PBMC tissue analysis (Fig 5). The unbiased set of top MRs for the individual subclusters include several functionally relevant genes that have been widely studied in relation to T-cell activity stratification. For example, cluster 1 presents highest activity of TCF7, a known master regulator of stem-like CD8 T cells^41,42^, as well as of CCR7, a regulator of T cell exit from peripheral tissues into lymphatic circulation^43^. Top Cluster 2 MRs include GZMA, a marker of activated cytotoxic CD8 T cells^44^, as well as CD160^45^, EOMES^46^, and FOSL2^47^, all associated in literature with CD8 T-cell activation. Cluster 3 is an unambiguous collection of B cells, with high differential activity of several canonical B cell markers (CD40, CD79A, CD24)^48,49^. Cluster 4 is positive for CXCR6, expressed on terminally differentiated effector CD4 T cells^50^. Finally, cluster 5 is positive for TNFRSF4 (OX40), a T cell immune checkpoint molecule actively being pursued as an immunotherapy target^51^ as well as IL2RA, induced widely among CD4 T cells in response to stimuli and constitutively expressed in regulatory T cells (Tregs)^52^. Critically, none of these markers appear differentially expressed within these clusters, due to gene dropout.

**Figure 5:**
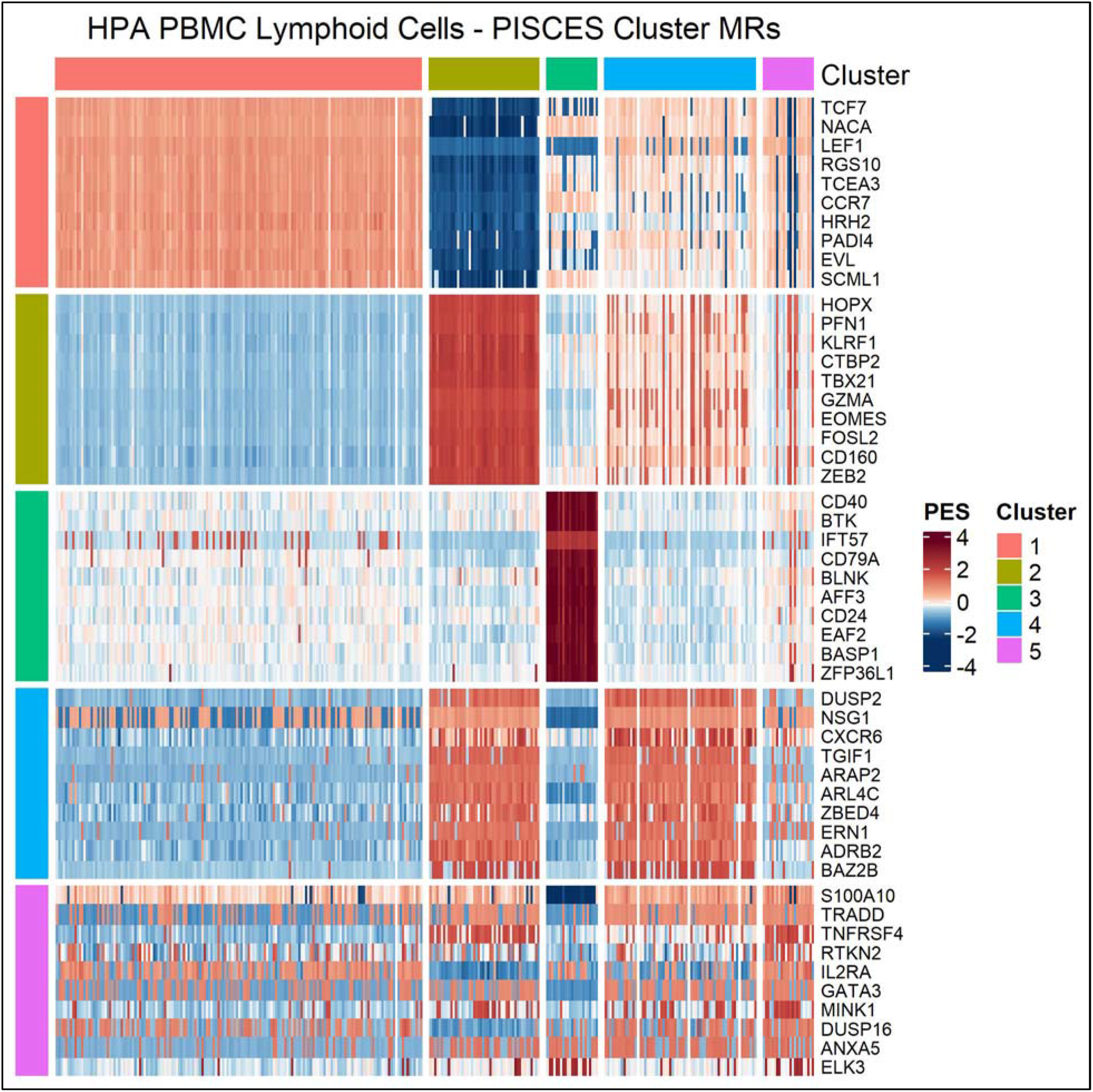
Hierarchical analysis of Lymphoid cells in HPA PBMC tissue Top-10 cluster-specific master regulators identified by PISCES clusters of Lymphoid cells in PBMC. Cluster 1 is identified as stem-like CD8+ T-cells, Cluster 2 as activated CD8+ T-cells, Cluster 3 as B-cells, Cluster 4 as CD4+ T-cells, and Cluster 5 as T-regs. Similar analysis was performed for all lineage-groups within across all tissues of the HPA.

## Discussion

PISCES provides a comprehensive, community-available, and highly customizable framework for the systematic assessment of protein activity and for the identification of molecular determinants and markers of single cell state. Specifically, we define protein activity as the protein-target mediated contribution of a protein to establish the transcriptional state of a cell.

Our study shows that the reproducibility of individual protein activity, as assessed by PISCES, significantly outperforms reproducibility of individual gene expression as the number of mRNA reads (UMI/cell) decrease. This addresses a core limitation intrinsic to the analysis of single-cell RNA-seq data. Furthermore, our analyses show that protein activity, as measured by PISCES, recapitulates experimental single cell protein abundance measurements—specifically, based on CITE-Seq antibody-based assays—again outperforming gene expression analyses. Finally, analysis of the Human Protein Atlas—which includes a comprehensive repertoire of single cell profiles from 26 human tissues—confirms that protein activity-based cluster and Master Regulator analyses can capture meaningful biological structure that would escape detection by gene expression-based analyses, as also supported by independent biological evidence.

PISCES applications far exceed the confines of what has been shown in this manuscript. Specifically, by allowing single cells to be analyzed quantitatively akin to bulk tissue, they allow expansion of a number of previously published VIPER-based methodologies, from the generation of machine learning-based classifiers to study outcome or drug resistance, to the identification of Master Regulator proteins, and single-cell-specific drug sensitivity prediction using the VIPER-based OncoTarget and OncoTreat algorithms. In addition, the growing popularity of single-nucleus and spatial transcriptomics RNAseq profiles, where gene dropout or low-gene repertoire availability is even further exacerbated, are even more likely to highlight the advantages of this platform; indeed single-nucleus profiles often report ≤ 2,000 UMI/cell^53^, and spatial transcriptomics and produce as few as 100 UMI/cell^54^.

In particular, an especially relevant conclusion of our studies is that, when high-quality networks are available, reproducible protein activity profiles can be generated with UMI/cell in these ranges, as low as 1,000 and potentially even lower. Since network architecture is lineage specific but not dataset specific, as the number of high-quality single cell tissue datasets increases, high quality networks can be generated using a variety of different approaches, for instance by consensus-based integration of networks generated by multiple dataset from lineage related subpopulations or even including networks from relatively pure bulk tissues in the metaVIPER2 network integration step.

Potential limitations of PISCES are largely centered on network generation. In most cases, network generation from single-cell data requires at least 250 cells, and more cells are typically required from low-depth data. This may lead to situations where it is not possible to generate networks for niche cell populations. MetaVIPER2 can integrate protein activity measurements using networks from similar cell types, but these results may not be as robust as using cell lineage-specific networks and there may still be scenarios where no networks are available for a closely related cell types. Generating networks is also computationally challenging due to the number of subnetworks required. We aim to address both of these limitations through the generation of network library from the HPA, providing researchers with a ready-made repository of high-quality networks covering a wide range of cell types across multiple tissues.

PISCES is made broadly available to the community from the Califano lab GitHub (https://github.com/califano-lab/PISCES). The repository of HPA networks and tissue-cluster-specific master regulator analyses can likewise be found on the Califano lab Dropbox, freely available for academic use (https://www.dropbox.com/sh/pc87euhnjrzess4/AAAuptJRAro-XDOjJWabrqDca?dl=0). Finally, all analysis code can be found in the PISCES validation repository of the Califano lab GitHub (https://github.com/califano-lab/PISCES_validation).

## Acknowledgements

This work was supported by an NCI Outstanding Investigator Award (R35CA197745), the NIH Instrumentation grants (S10OD012351 and S10 OD021764) and the Chan Zuckerberg Initiative CZI) to A.C.. Also, this research was funded in part through the NCI Cancer Center Support Grant (P30 CA013696).

## Online Methods

### Quality Control

As a pre-processing step, low quality cells and genes lacking enough data to be useful are removed from the analysis. Cell quality is determined by three factors; read depth, number of detected genes, and mitochondrial gene percentages. Samples with too many or too few reads are likely sequencing errors (doublets or empty droplets); cells with too few detected genes are also likely sequencing errors or dying cells; and high mitochondrial gene percentage is indicative of cell stress or damage.

### Metacell Generation

Metacells are generated through a k-nearest-neighbors approach. A distance matrix is first calculated in gene expression space – typically a Euclidean distance metric based on the first 10-30 principle components of either the Log2(CPM + 1) normalized counts or from the SCT matrix generated by Seurat is used. Next, nearest neighbors are identified, and the raw counts for each cell are summed with those from its k nearest neighbors. The number of neighbors should be set such that the average depth of the resulting metacells is > 10,000 UMIs / cell. Metacell counts are then Log2(CPM + 1) normalized. Finally, we recommend subsetting the final metacell matrix to between 250 and 500 metacells. Because ARACNe assumes independence of samples, overly dense metacell generation can confound network generation. For a given number of cells *n*, metacell size *k*, and subset size *s*, each cell will be included on average 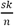 times in a metacell matrix. Thus, by using a smaller subset *s*, the overlap probability can be reduced, allowing independence of samples to largely be maintained and for robust networks to be generated.

### ARACNe3 Network Generation

Networks are generated using ARACNe3, the latest implementation of the ARACNe algorithm. Briefly, ARACNe3 takes in a list of putative regulators and identifies a set of targets (i.e. a regulon) based on the mutual information between the expression of the regulator and the target. Trios in the network are pruned based on the principle of maximum entropy to avoid local cycles. Typically, multiple networks are generated from subsamples of the original gene expression profile, with 1 – 1/e percent of the data represented in each subsample. These subsample networks are then integrated into a final regulatory network, consisting of the regulons for each of the putative regulators for which there were a sufficient number of targets.

There are four primary parameters to consider when running ARANCe3 on single-cell data; the depth of the gene expression profile; the number of cells; the FDR of the ARACNe3 null model; and the number of subnetworks used. We perform an exploration of this parameter space in Figure 2B-D. All networks were generated using a script specific to our HPC architecture, available on the PISCES github. Note that this script will require modifications for different HPC systems.

### MatrixNaRnEA

Matrix NaRnEA adapts the original NaRnEA algorithm to be applied to all samples in an input GES and all regulons in an input network simultaneously. The ‘matrix_narnea’ package takes in a gene expression signature matrix (genes in rows, samples in columns) and an appropriately parametrized regulatory network list. The network list should be a named list, where the names are regulators and the items are regulons; the regulons themselves should be a list with association weight (‘aw’) and association mode (‘am’) elements, each of which is a named list of targets with the corresponding parameter.

Hereafter, we refer to the input gene expression signature matrix as *G*, and elements of *G* as *g*_*ij*_. The number of genes in the signature is given as *h*, the number of samples in the signature as *n*, and the number of regulons in the network as *q*. Note also that ‘×’ refers to matrix multiplication, while ‘⋅’ refers to element-wise multiplication. Subscripts will point to targets *t*, regulators *r*, and samples in the gene expression signature *x*. The targets of regulator *j* are referred to as *regj*. In general, derivations will start with the single-regulon, single-sample definition before expanding to the vectorized equations. For a full derivation of NaRnEA, please see the NaRnEA manuscript^27^.

MatrixNaRnEA starts by defining the following values:

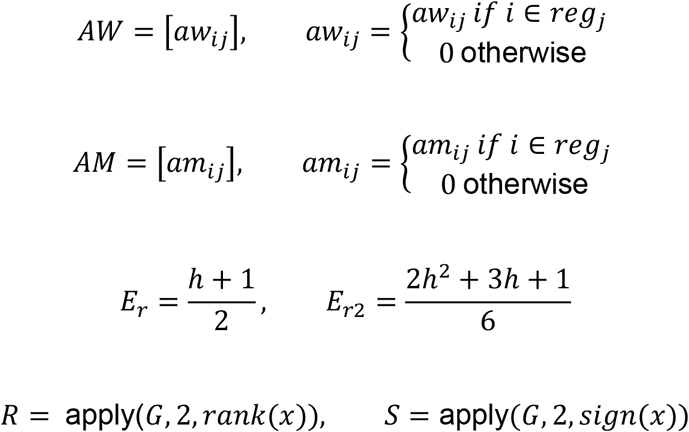

The Directed Enrichment Score matrix *D* and it’s corresponding expectation, variance, and normalized enrichment score are calculated as follows:

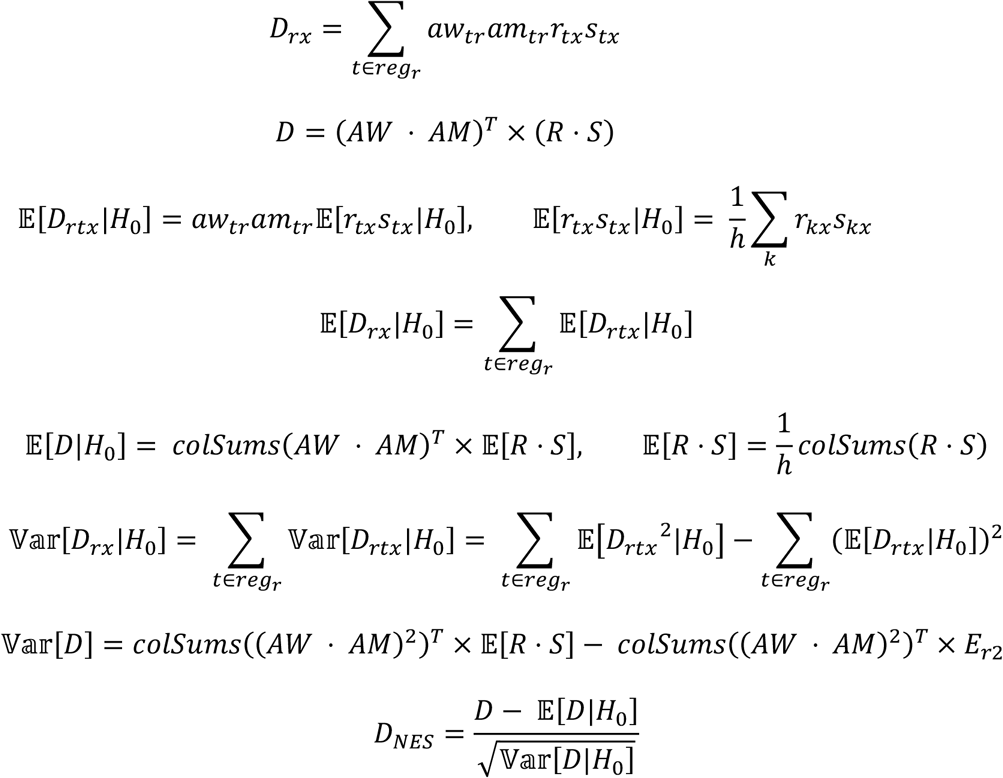

The Undirected Enrichment Score matrix U and it’s corresponding expectation, variance, and normalized enrichment score are calculated as follows:

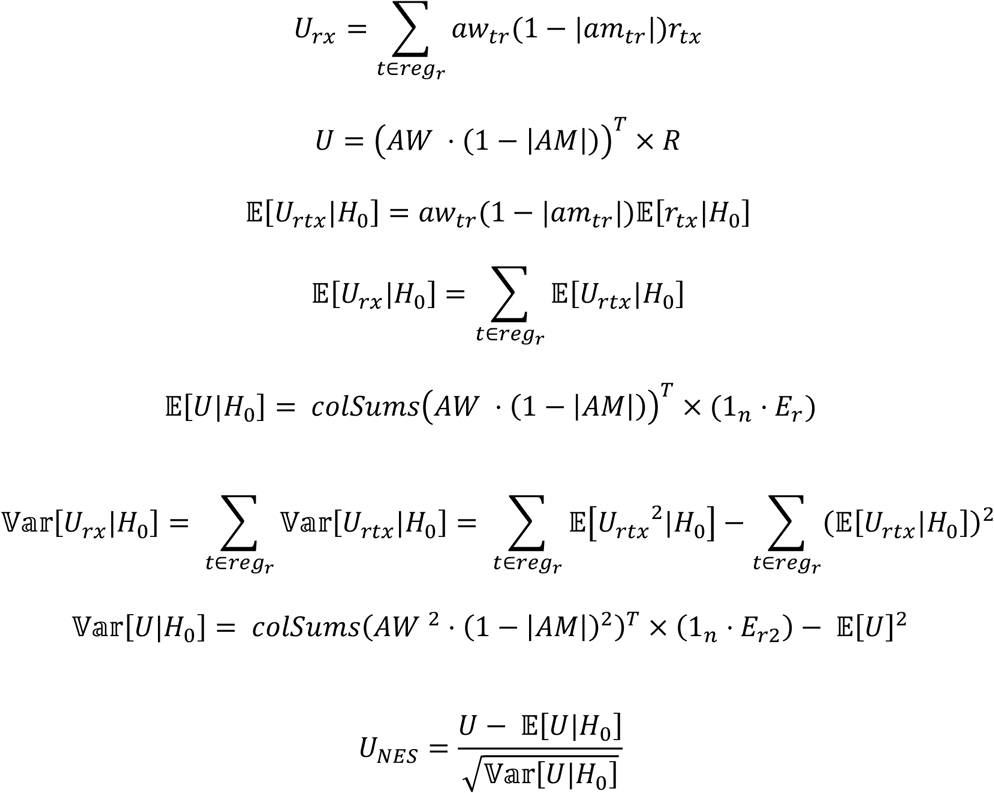

We then calculate the covariance between the Undirected and Directed Enrichment Score as follows:

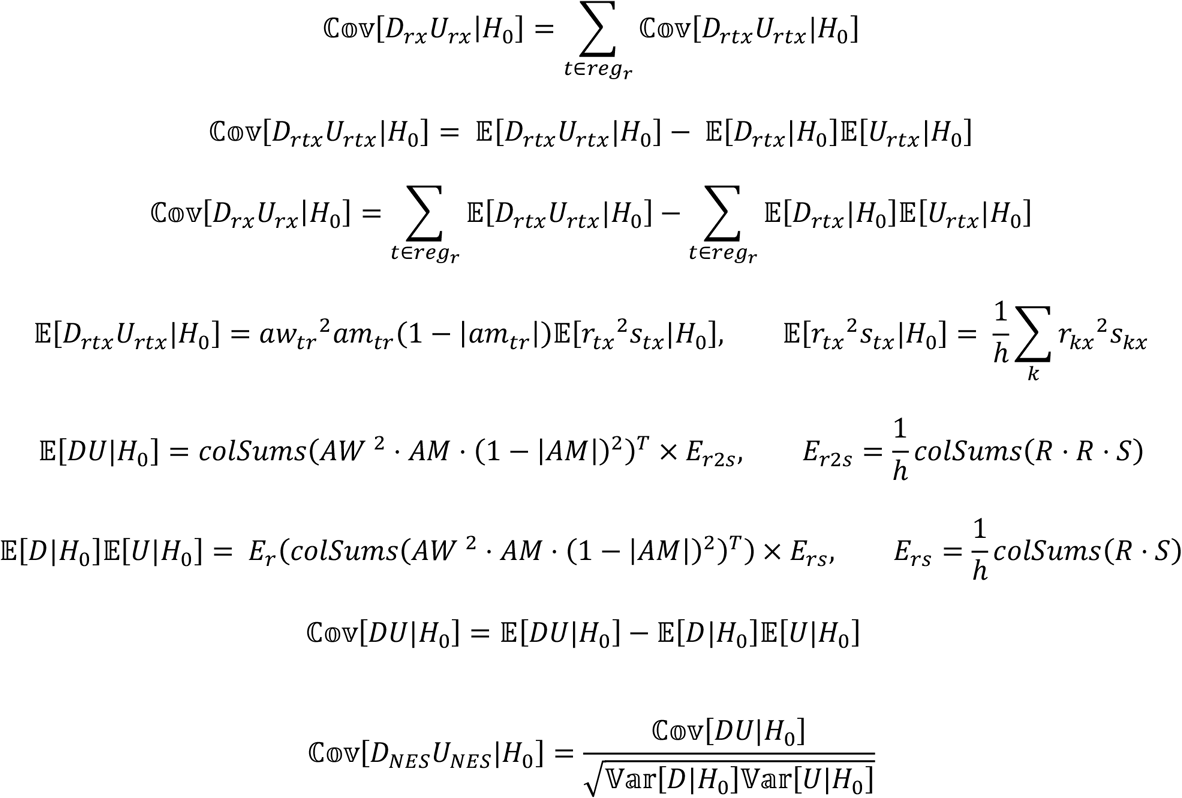

We can now combine the Directed and Undirected Normalized Enrichment Scores into a final NES value:

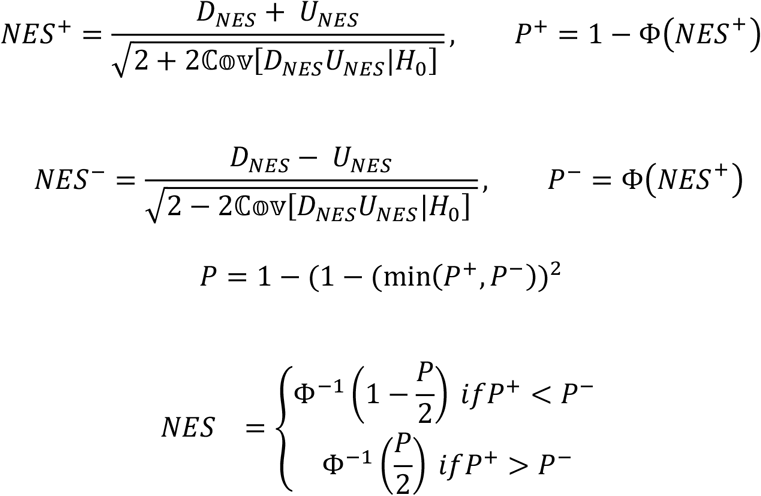

Finally, we calculate the proportional enrichment score (PES) by first calculating a maximum theoretical value for each regulon. For each regulon, the target genes are sorted by the strength of the association mode. A dummy signature is created such that the nth most strongly regulated gene in the regulon is the nth most differentially expressed gene in the signature. Additionally, the sign of the signature is taken to be the same as that of the association mode. A DES and PES is calculated from this dummy signature, normalized by the appropriate means and variances that were previous calculated, then integrated into a final theoretical maximum NES. The PES is then calculated by dividing the actual NES in each regulon for each sample by the theoretical maximum NES for the appropriate regulon.

Note that this is slightly divergent from the approach used with the original NaRnEA algorithm – here the maximum values are tuned to the signature as a hole as opposed to each sample specifically, and no regard is paid to the underlying distribution of the sign of differentially expressed gene. This will, in general, lead to numerically smaller PES values, though the change is so minimal as to be irrelevant. For a full derivation of the original PES values, see the NaRnEA manuscript.

### MetaNaRnEA Analysis

MetaNaRnEA integrates NaRnEA results from multiple networks into a final NaRnEA object. Under the null hypothesis of NaRnEA – a given regulator shows no differential activity – we expect networks that do not accurately recapitulate the biology of the given sample to show no enrichment, as regulons with randomly selected targets have been demonstrated to show no differential activity. We leverage this knowledge to appropriately identify and weight networks for integration. For each protein in each sample, the network that returns the largest PES is taken as the network that best matches the sample in question. This is repeated for each regulator shared across the networks, generating a count of the number of times each network provided the optimally matched regulon. These counts then serve as weights for integrating NaRnEA values; PES scores are combined via a weighted average, while NES scores are combined using weighted Stouffer integration. For regulators that are not shared across all networks used for the analysis, weights are re-normalized over the networks in which the regulon is present.

This functionality is implemented using matrix arithmetic to speed computation but can still be slow to run for large number of cells. An unweighted option is also available for particularly large datasets, but we cannot guarantee the quality of these results. This functionality is included in the ‘meta_narnea’ function in PISCES.

### Resolution-Optimized Louvain Clustering Algorithm

The default clustering method implemented in Seurat is Partitioning Around Medioids (PAM). However, for large datasets aggregating hundreds of thousands of single-cells, PAM is computationally slow, requiring more computational power than is available to the average user and computation of pairwise distance matrices exceeding the vector size limit in R. To overcome this, we utilize the ‘igraph’ package in R to implement silhouette-score optimized Louvain clustering. A k-nearest-neighbor graph is created from a given distance matrix, with k being a free parameter that is optimized over. By default, the ‘louvain_k’ function will test values of k between 5 and 50 at intervals of 5, returning both a vector of all clustering solutions and a single optimal solution based on the highest silhouette score.

### Multi-Way K-Means Clustering Algorithm

In addition to PAM and Louvain with Resolution Optimization, PISCES further implements a Multi-Way K-Means Clustering approach. Transitioning populations, such as in a differentiation pathway, are extremely common, and such relationships will not be accurately characterized by a discrete clustering scheme. To handle this set of problems, we adapted the Multiway K-Means algorithm for use in biological settings, where samples can be thought of as linear combinations of related phenotypes rather than simply belonging to totally distinct populations. Originally developed for clustering speciating microbiome populations, Multiway K-Means technique has two primary advantages. First, it more accurately captures cluster center (in biological terms, a representative phenotype) for each population endpoint. Second, it places cells along a trajectory between cluster centers, providing a more accurate representation of cell state and allowing for additional inferences into the drivers of transitional populations.

### Master Regulator Analysis

PISCES implements several options for master regulator analysis between populations. The default option is to run a Kruskal-Wallis test between the PES values of each protein and the vector of cluster membership. P-values are corrected by the method of Benjamini and Hochberg, and group-specific master regulators are sorted based on the group residual of each protein. We present both the positive (activated) and negative (deactivated) master regulators as separate lists. This procedure is performed by the ‘kw_cluster_mrs’ function. For all analyses performed in this manuscript, this methodology was used to identify cluster-specific master regulators.

Master regulators can also be identified by the integration of the protein activity signature across a group. NES values across a group or cluster are Stouffer integrated, and the integrated NES for each protein is converted to a p-value. These p-values are then corrected by the method of Benjamini and Hochberg, and the final list of positive and negative group-specific MRs are sorted by the mean of the average PES across the group. This methodology is appropriate when comparing an entire sample against an external reference, for example. This procedure is performed by the ‘stouffer_cluster_mrs’ function.

Finally, we also provide a machine-learning flavored method for inferring master regulators based on Cohen’s kappa coefficient^55^. For each protein in each group, a single-feature decision tree is trained to best differentiate the group in question versus the rest of the population. Proteins within each group are then sorted by the ability of their classifier to best distinguish the group as measured by Cohen’s kappa coefficient. Master regulators are finally signed by the average in-group PES, and the procedure is repeated for each group in the dataset. This method is anecdotally strong at identifying the most differentiation features for single groups. This procedure is performed by the ‘kappa_cluster_mrs’ function.

### Read Subsampling Procedure

Reads were subsampled in a sample-specific manner to generate depth-controlled gene expression profiles in a two step stochastic sampling process. First, cell-specific depths were modeled as a multinomial distribution, with each cell having an equal probability of receiving a read and a total number of reads equal to the number of cells times the target average depth being drawn. Next, the reads within each cell were modeled as a cell-specific multinomial, with the previously drawn cell-depth as the total number of draws and the frequency of each gene in the full-depth data as the gene-specific probability. As a final step, any rows with no genes were removed from the final read-subsampled gene expression profile. This procedure was used for both the technical validation and for the read subsampling of the Sims CITE-Seq dataset.

### Selecting optimal ARACNe3 parameters

SNU-16 data was stochastically read-subsampled to depths 2,000, 5,000, and 10,000 UMIs / cell. ARACNe3 regulatory networks were then generated from each of these depths varying three parameters; the number of samples in the starting data (testing 250, 500, and 1000 samples); the number of subnetworks used (testing 5, 10, 25, and 50); and finally the FDR used in the mutual information null model (testing 0.05, 0.25, and 0.5). For each possible parameter configuration, we created three ARACNe3 networks. Additionally, we computed profile-level gene expression signatures for each read-subsampled profile against CCLE and generated a NaRnEA signature for each network parameter configuration.

To measure the effect of each parameter combination on reproducibility, we generated an ARACNe3 network from the full-depth SNU-16 data and computed a profile-level signature against appropriately read-subsampled CCLE data. We then computed NaRnEA enrichment of the full depth signature using the full depth network, taking this as a “gold standard” protein activity signature. Network parameter configurations were then evaluated based on the enrichment of this “gold standard” signature in the network-generated signatures as inferred by NaRnEA (Figures 2B-C).

This same reproducibility assessment procedure was applied to the network generated with metacells from the 2,000 and 5,000 UMI datasets.

### Features Recovered by Protein Activity

SNU-16 data was read-subsampled to depths between 1,000 and 10,000 UMIs / cell. Genes with fewer than 6 reads – equivalent to 0.1% times the number of cells (6,157) – were filtered as lacking enough data. Single-cell gene expression signatures were generated against appropriately read-subsampled CCLE. Protein activity was inferred using MatrixNaRnEA with the full-depth SNU-16 network. Proteins for which we aimed to compute activity were regarded as the feature set of interest. Features were counted based on the overlap of genes in the given gene expression profile or NaRnEA signature and the feature set of interest (Supplementary Table 4). Significant features were regarded as those that produced an absolute NES value equivalent to a Bonferroni p-value of 0.05 in the 95^th^ percentile of cells. We used the 95^th^ percentile rather than the maximum to avoid the influence of outlier cells. This procedure was repeated for both the read-subsampled gene expression signature and the NaRnEA signature generated from the read-subsampled GES and the full-depth network, and the set of significant features was selected as the overlap of these features with the feature set of interest (Supplementary Table 5).

### Normalization and Scaling Parameters Tested

We tested three different methods of depth-normalizing the input counts. First, we applied the widely used counts-per-million (CPM) approach, normalizing the depth of all cells to a size factor of one million before log2 transforming the counts. Second, we utilized the single cell transformation (SCT) method from Satija et. al., which transforms data to Pearson residuals derived from a regularized negative binomial regression^7^. Finally, we applied the two-step proportional fitting approach recommended by Booeshagi et. al., which has been reported as the most effective method for removing the effect of depth on downstream analyses^56^. We refer to these three methods as “CPM”, “SCT”, and “PFLPF” respectively.

To generate sample-specific gene expression signatures (GES), we tested two different methods of comparing depth-normalized counts for a given gene in one sample versus the rest of the population. The first was a simple internal scaling, subtracting the mean normalized count and dividing by the standard deviation. Second, we utilized the empirical cumulative distribution function (ECDF) for each gene, first transforming the depth-normalized expression into a quantile with the ‘ecdf’ function, then transforming the result into a z-score using ‘qnorm’. These two methods are referred to as “scale” and “ecdf” respectively.

Additionally, we tested the effectiveness of including a Jeffrey’s Prior in our signature generation step^57^. For compositional data, this is equivalent to the inclusion of a pseudo-count of 0.5 for each gene. Results from these signature include a “pseudo” clause in the name.

We find that the majority of the possible combinations of these methods perform well in terms of generating protein activity measurements that recapitulate the antibody abundance levels generated from CITE-Seq. In general, “CPM” and “scale” lead to the best results, with “sct” typically being the next best option. Interestingly, the inclusion of pseudo-counts, though well justified from the perspective of Bayesian statistics, typically reduced rather than improved performance. For a full view of the performance of these options, see Supplementary Table 2.

### Network Generation Groups Tested

We tested three potential methods for segregating cells at the gene expression prior to network generation; an unsupervised approach based on Seurat clustering; and two supervised approaches based on SingleR, both at the cell-type and lineage level. We found that grouping networks into lineage-specific groups (i.e. combining T-Cell and B-Cell populations into a single Lymphoid group rather than making separate networks) generated better results as measured by the reproducibility of the resulting protein activity signatures with antibody abundance measurements from CITE-Seq. This is likely due to these groups of cells sharing regulatory architecture due to being from the same lineage. As such, we generally recommend performing analysis to group cells into broad, lineage based groups (i.e. lymphoid, myeloid, epithelial, fibroblast, etc.) for network generation. A typical approach would be to perform unsupervised clustering, then analyze marker genes or SingleR results to map broad major lineages or cell types onto these clusters, regrouping as appropriate. We note when this approach is not viable, networks simply generated simply from the unsupervised clusters will still generate robust protein activity signatures, however.

### NaRnEA Integration Methods Tested

In addition to the MetaNaRnEA integration method described previously, we tested an alternative, more supervised approach to combining NaRnEA inferences from multiple networks. Network Match NaRnEA (implemented in the function ‘net_match_narnea’) requires an additional vector indicating which networks match each sample in the given gene expression signatures. For samples with an explicitly matched network, only that network will be used for protein activity measurement. Those samples without explicitly matched networks will have protein activity measurements generated using MetaNaRnEA. This contrasts with the MetaNaRnEA approach, where all networks are considered for all samples and the network weights are derived based on the best matching network as determined by the data.

We find that in general, MetaNaRnEA outperforms Network Match NaRnEA for generating protein activity measurements that more accurately recapitulate antibody abundance levels as measured by CITE-Seq. Network Match NaRnEA results still do a reasonable job and typically outperform gene expression methods, but we generally recommend the MetaNaRnEA approach. For a full view of the performance of these options, see Supplementary Table 2.

### PBMC Regulatory Network Generation

ARACNe3 networks were generated on the C2B2 HPC at Columbia University. Networks for CITE-Seq analysis were generated were the most generalizable parameters; null model FDR of 0.25, 50 subnetworks, and 500-1,000 samples per network generation group. Three separate runs were performed for each network, one for each of TFs and COTFs, signaling proteins and surface markers. The resulting networks were then concatenated into a final network used for subsequent analysis. The scripts and marker sets used for this analysis can be found in the on the Califano lab Github in the PISCES repository. Note that these scripts will likely need to be adapted to different HPCs. Additionally, further advances have since been made with regards to ARACNe3, and a C++ implementation that makes local runtimes potentially viable is available on the Califano lab GitHub (https://github.com/califano-lab/ARACNe3). Results from this new implementation will not be identical due to the stochastic nature of subsampling, but will produce concordant results with the analyses described previously.

### Correlation Analysis

The change in correlation with antibody abundance between gene expression and protein activity was analyzed using the ‘cocor’ package in R. A set of comparison genes was identified by taking the intersection between genes that passed quality control, proteins for which a regulon could be inferred and protein activity measured, and genes for which antibody abundance was available. Correlations were modeled as dependent and calculated by first rank-transforming the vector of counts / PES values, then computing correlation with the ‘cor’ function in R, generating spearman correlation. We used the ‘dunn1969’ value from CoCor, which leverage’s Dunn and Clark’s z test statistic^58^.

### PBMC Data Generation

CITE-Seq data from the Sims lab was generated as described in Conde et al.^59^.

### PISCES Visualizations

Heatmaps were generated using the ‘complexHeatmap’ package in R^60^. Flowcharts were generated using Microsoft PowerPoint. All other figures were generated using the ‘ggplot2’ packages in R^61^.

### HPA Analysis

Tissues were analyzed using the parameters identified as optimal by our validation experiments. To reduce memory load an computational time, several tissues were subset to 10,000 or 15,000 cells. A full summary of tissue analyses with QC parameters, network details, and subsetting parameters are included in the HPA Analysis Dropbox.

### Data Availability

PBMC CITE-Seq data from Seurat are available in their vignette (https://satijalab.org/seurat/articles/multimodal_vignette.html). PBMC Cite-Seq data generated by the Sims lab is available on their GitHub (https://github.com/simslab/pbmc_citeseq). SNU-16 data utilized for technical validation can be found on the Califano lab GitHub (https://www.dropbox.com/sh/u42hzb6ad3tg9qr/AABRvsdW0dlVNzJSIl35Cx5Ua?dl=0).

### Code Availability

Code used for validation analyses can be found on the Califano lab GitHub (https://github.com/califano-lab/PISCES_validation). Code for the PISCES analysis of the HPA can be found on the Califano lab Dropbox along with all results and logs of HPC commands (https://www.dropbox.com/home/Califano%20Lab%20Shared/PISCES_HPA/code).

---

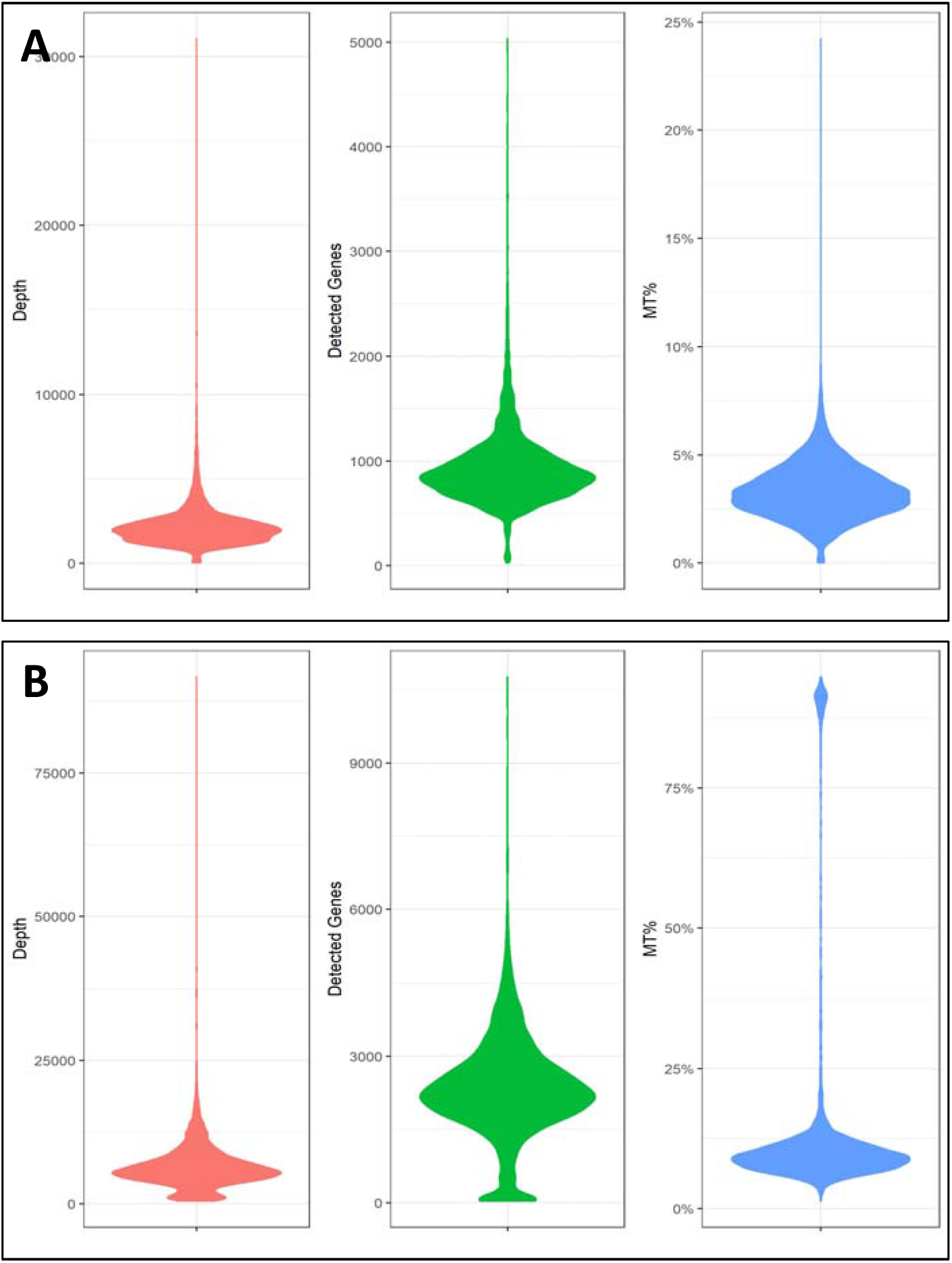

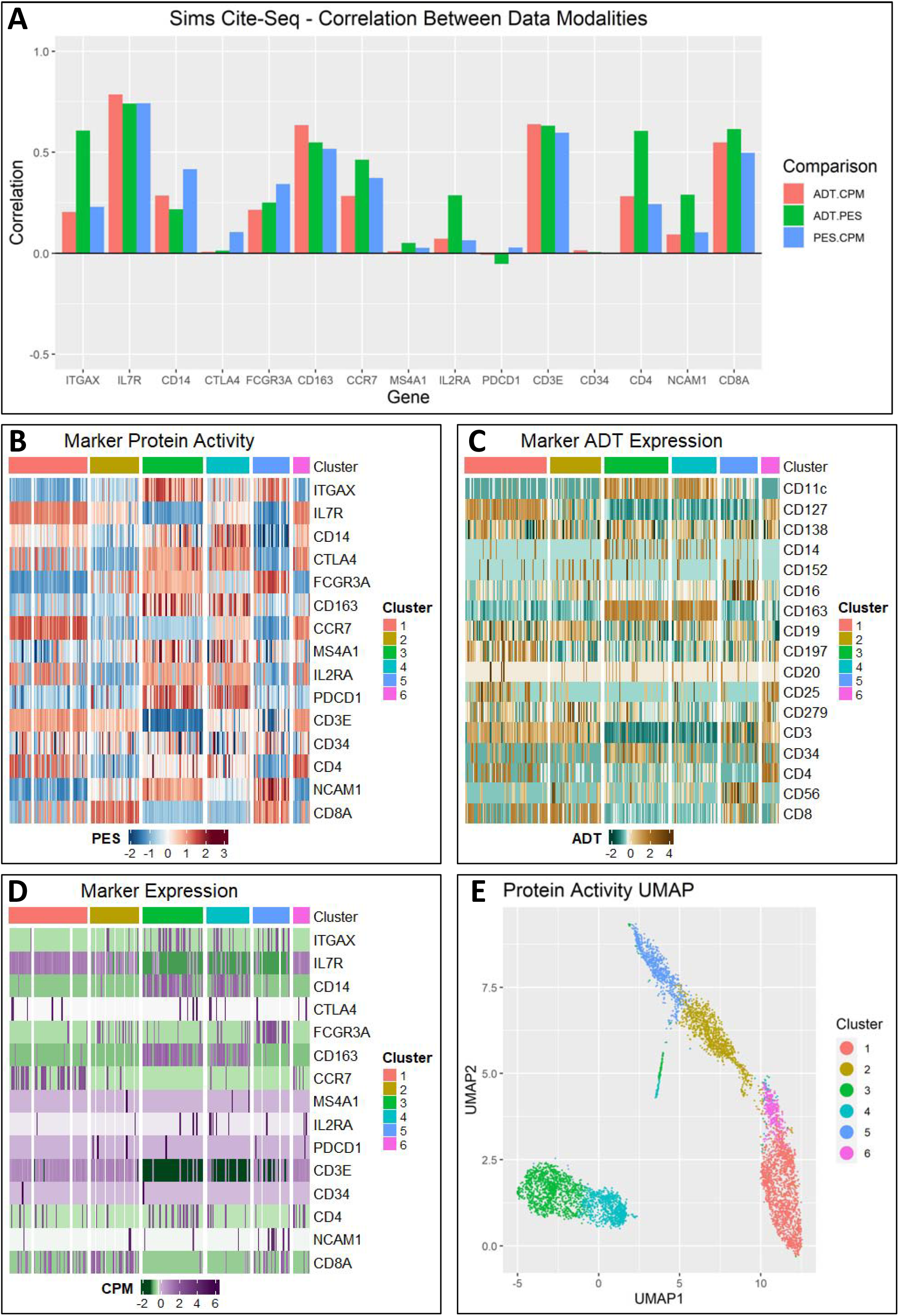

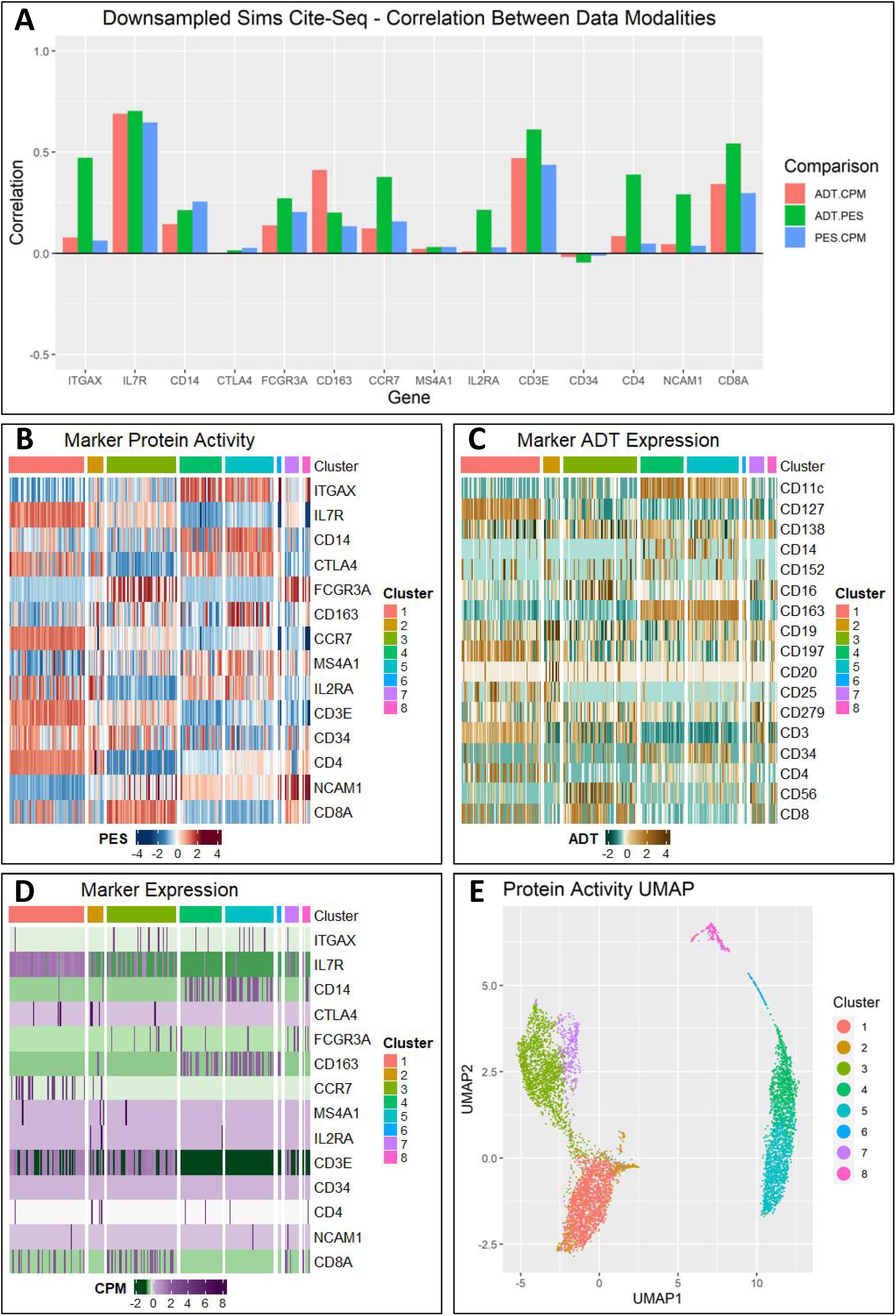

**Supplementary Table 1:** QC Thresholds for validation datasets

| Dataset | Max MT% | Min. UMI Depth | Max UMI Depth | Min. Unique Genes | Max Unique Genes |
| --- | --- | --- | --- | --- | --- |
| Seurat CITE-Seq | 0.1 | 1000 | 10000 | 725 | 3000 |
| Sims CITE-Seq | 0.25 | 4000 | 37500 | 1000 | 7500 |

**Supplementary Table 2:** Full parameter exploration for PISCES, sorted by improved concordance w/ antibody abundance relative to gene expression

| Signature | Network | Method | Zscore | logp | ADT.CPM | ADT.PES | num.genes |
| --- | --- | --- | --- | --- | --- | --- | --- |
| cpm-scale | sr-lineage | meta-narnea | 66.66 | -2227.03 | 0.14 | 0.41 | 11 |
| pflpf-scale | sr-lineage | meta-narnea | 65.32 | -2138.65 | 0.15 | 0.41 | 11 |
| pflpf-scale | sr-lineage | net-match-narnea | 52.48 | -1382.19 | 0.14 | 0.34 | 11 |
| cpm-scale | sr-lineage | net-match-narnea | 52.24 | -1369.55 | 0.14 | 0.34 | 11 |
| cpm-ecdf | sr-lineage | meta-narnea | 46.96 | -1107.50 | 0.14 | 0.32 | 11 |
| sct-full | sr-lineage | meta-narnea | 46.60 | -1090.62 | 0.14 | 0.33 | 11 |
| sct | sr-lineage | meta-narnea | 46.00 | -1062.95 | 0.14 | 0.33 | 11 |
| pflpf-ecdf | sr-lineage | meta-narnea | 45.64 | -1046.07 | 0.14 | 0.32 | 11 |
| sct | sr-lineage | net-match-narnea | 39.22 | -773.75 | 0.14 | 0.30 | 11 |
| cpm-pseudo-ecdf | sr-lineage | net-match-narnea | 38.69 | -752.85 | 0.14 | 0.31 | 11 |
| pflpf-pseudo-ecdf | sr-lineage | net-match-narnea | 35.78 | -644.76 | 0.14 | 0.30 | 11 |
| sct-full | sr-lineage | net-match-narnea | 35.25 | -625.71 | 0.14 | 0.28 | 11 |
| cpm-pseudo-ecdf | sr-lineage | meta-narnea | 34.93 | -614.42 | 0.14 | 0.29 | 11 |
| pflpf-pseudo-ecdf | sr-lineage | meta-narnea | 33.50 | -565.67 | 0.14 | 0.28 | 11 |
| pflpf-scale | sr-ct | meta-narnea | 29.88 | -450.72 | 0.14 | 0.27 | 11 |
| sct | sr-ct | meta-narnea | 24.68 | -308.61 | 0.15 | 0.25 | 11 |
| cpm-scale | sr-ct | meta-narnea | 24.58 | -306.16 | 0.14 | 0.25 | 11 |
| cpm-pseudo-ecdf | sr-ct | meta-narnea | 21.13 | -227.12 | 0.14 | 0.23 | 11 |
| pflpf-pseudo-ecdf | sr-ct | meta-narnea | 20.32 | -210.41 | 0.14 | 0.23 | 11 |
| sct-full | sr-ct | meta-narnea | 20.27 | -209.35 | 0.14 | 0.23 | 11 |
| pflpf-ecdf | sr-lineage | net-match-narnea | 18.38 | -172.70 | 0.14 | 0.22 | 11 |
| pflpf-ecdf | sr-ct | meta-narnea | 17.60 | -158.65 | 0.14 | 0.20 | 11 |
| cpm-ecdf | sr-ct | meta-narnea | 12.57 | -82.41 | 0.14 | 0.18 | 11 |
| cpm-ecdf | sr-lineage | net-match-narnea | 10.75 | -61.08 | 0.15 | 0.19 | 11 |
| sct | seurat-clust | net-match-narnea | 9.97 | -52.97 | 0.14 | 0.17 | 11 |
| sct | sr-ct | net-match-narnea | 6.24 | -22.27 | 0.14 | 0.16 | 11 |
| cpm-pseudo-ecdf | sr-ct | net-match-narnea | 6.04 | -21.00 | 0.15 | 0.18 | 11 |
| pflpf-pseudo-ecdf | sr-ct | net-match-narnea | 5.45 | -17.52 | 0.14 | 0.17 | 11 |
| sct | seurat-clust | meta-narnea | 4.36 | -11.94 | 0.14 | 0.14 | 11 |
| cpm-scale | seurat-clust | meta-narnea | 4.36 | -11.93 | 0.14 | 0.14 | 11 |
| pflpf-scale | seurat-clust | meta-narnea | 4.23 | -11.34 | 0.14 | 0.14 | 11 |
| cpm-pseudo-ecdf | seurat-clust | meta-narnea | 0.07 | -0.75 | 0.14 | 0.13 | 11 |
| pflpf-pseudo-ecdf | seurat-clust | meta-narnea | -1.93 | -0.03 | 0.15 | 0.13 | 11 |
| cpm-scale | sr-ct | net-match-narnea | -2.80 | 0.00 | 0.14 | 0.11 | 11 |
| pflpf-scale | sr-ct | net-match-narnea | -3.50 | 0.00 | 0.15 | 0.12 | 11 |
| sct-full | seurat-clust | meta-narnea | -4.15 | 0.00 | 0.14 | 0.11 | 11 |
| sct-full | sr-ct | net-match-narnea | -7.73 | 0.00 | 0.14 | 0.09 | 11 |
| pflpf-scale | seurat-clust | net-match-narnea | -11.46 | 0.00 | 0.15 | 0.08 | 11 |
| sct-full | seurat-clust | net-match-narnea | -11.81 | 0.00 | 0.14 | 0.07 | 11 |
| cpm-scale | seurat-clust | net-match-narnea | -12.42 | 0.00 | 0.14 | 0.07 | 11 |
| pflpf-pseudo-ecdf | seurat-clust | net-match-narnea | -13.48 | 0.00 | 0.14 | 0.07 | 11 |
| cpm-pseudo-ecdf | seurat-clust | net-match-narnea | -15.12 | 0.00 | 0.14 | 0.07 | 11 |
| cpm-ecdf | seurat-clust | meta-narnea | -17.88 | 0.00 | 0.13 | 0.04 | 11 |
| pflpf-ecdf | seurat-clust | meta-narnea | -24.35 | 0.00 | 0.15 | 0.02 | 11 |
| cpm-ecdf | sr-ct | net-match-narnea | -32.92 | 0.00 | 0.14 | -0.03 | 11 |
| pflpf-ecdf | sr-ct | net-match-narnea | -33.79 | 0.00 | 0.15 | -0.03 | 11 |
| pflpf-ecdf | seurat-clust | net-match-narnea | -38.18 | 0.00 | 0.14 | -0.06 | 11 |
| cpm-ecdf | seurat-clust | net-match-narnea | -38.39 | 0.00 | 0.14 | -0.06 | 11 |

**Supplementary Table 3:** Full parameter exploration for PISCES clustering options, sorted by concordance with antibody clustering as measured by silhouette score

| Signature | Network | Method | Distance | ADT.silhouette | KW.logp |
| --- | --- | --- | --- | --- | --- |
| cpm-scale | sr-lineage | meta-narnea | pca-euc | 0.040919083 | -21709.97254 |
| cpm-scale | sr-lineage | meta-narnea | pca-cor | 0.003779169 | -20814.88915 |
| cpm-scale | sr-lineage | meta-narnea | euc | 0.285601024 | -22392.65044 |
| cpm-scale | sr-lineage | meta-narnea | cor | 0.28159577 | -22300.88787 |
| pflpf-scale | sr-lineage | meta-narnea | pca-euc | 0.016207418 | -22175.10237 |
| pflpf-scale | sr-lineage | meta-narnea | pca-cor | 0.009438213 | -20993.61151 |
| pflpf-scale | sr-lineage | meta-narnea | euc | 0.205951538 | -22039.33063 |
| pflpf-scale | sr-lineage | meta-narnea | cor | 0.274881338 | -21177.32512 |
| pflpf-scale | sr-lineage | nm-narnea | pca-euc | 0.017509613 | -21477.67551 |
| pflpf-scale | sr-lineage | nm-narnea | pca-cor | -0.033937168 | -19716.91225 |
| pflpf-scale | sr-lineage | nm-narnea | euc | 0.317621048 | -19840.04433 |
| pflpf-scale | sr-lineage | nm-narnea | cor | 0.329545107 | -20143.35113 |
| cpm-scale | sr-lineage | nm-narnea | pca-euc | 0.067704502 | -21665.25334 |
| cpm-scale | sr-lineage | nm-narnea | pca-cor | -0.039320921 | -19767.58024 |
| cpm-scale | sr-lineage | nm-narnea | euc | 0.261356562 | -21105.65779 |
| cpm-scale | sr-lineage | nm-narnea | cor | 0.23696255 | -20797.63521 |
| cpm-ecdf | sr-lineage | meta-narnea | pca-euc | -0.073196934 | -20128.25037 |
| cpm-ecdf | sr-lineage | meta-narnea | pca-cor | -0.008260709 | -20643.43487 |
| cpm-ecdf | sr-lineage | meta-narnea | euc | -0.01699321 | -19635.92367 |
| cpm-ecdf | sr-lineage | meta-narnea | cor | 0.376919182 | -17635.41398 |
| sct-full | sr-lineage | meta-narnea | pca-euc | -0.053594427 | -21004.4133 |
| sct-full | sr-lineage | meta-narnea | pca-cor | -0.053056272 | -19549.76226 |
| sct-full | sr-lineage | meta-narnea | euc | 0.067858151 | -19644.16974 |
| sct-full | sr-lineage | meta-narnea | cor | 0.111709394 | -20294.55238 |
| sct | sr-lineage | meta-narnea | pca-euc | -0.076390752 | -21791.43659 |
| sct | sr-lineage | meta-narnea | pca-cor | -0.064249412 | -20387.10613 |
| sct | sr-lineage | meta-narnea | euc | 0.111382649 | -20629.9039 |
| sct | sr-lineage | meta-narnea | cor | 0.069231079 | -20507.34634 |
| pflpf-ecdf | sr-lineage | meta-narnea | pca-euc | -0.071060855 | -20939.38836 |
| pflpf-ecdf | sr-lineage | meta-narnea | pca-cor | -0.098765349 | -19926.78404 |
| pflpf-ecdf | sr-lineage | meta-narnea | euc | -0.018535997 | -19037.7013 |
| pflpf-ecdf | sr-lineage | meta-narnea | cor | 0.085552054 | -17627.84434 |
| sct | sr-lineage | nm-narnea | pca-euc | -0.015282559 | -20318.96623 |
| sct | sr-lineage | nm-narnea | pca-cor | -0.041978399 | -19527.70183 |
| sct | sr-lineage | nm-narnea | euc | 0.244965529 | -17985.85379 |
| sct | sr-lineage | nm-narnea | cor | 0.288922201 | -19875.04994 |
| cpm-pseudo-ecdf | sr-lineage | nm-narnea | pca-euc | -0.120180726 | -19842.57903 |
| cpm-pseudo-ecdf | sr-lineage | nm-narnea | pca-cor | -0.242361233 | -19083.37187 |
| cpm-pseudo-ecdf | sr-lineage | nm-narnea | euc | -0.123120127 | -19567.9057 |
| cpm-pseudo-ecdf | sr-lineage | nm-narnea | cor | -0.113880024 | -19340.93374 |

**Supplementary Table 4:** Feature count for protein activity (PISCES) and gene expression (GEP) at subsampled depths

| Depth | 1000 | 2000 | 3000 | 4000 | 5000 | 6000 | 7000 | 8000 | 9000 | 10000 |
| --- | --- | --- | --- | --- | --- | --- | --- | --- | --- | --- |
| NaRnEA | 5641 | 5641 | 5641 | 5641 | 5641 | 5641 | 5641 | 5641 | 5641 | 5641 |
| GEP | 4291 | 4520 | 4674 | 4755 | 4831 | 4893 | 4942 | 4982 | 5038 | 5064 |

**Supplementary Table 5:** Count of significant features for protein activity (PISCES) and gene expression (GEP) at subsampled depths

| Depth | 1000 | 2000 | 3000 | 4000 | 5000 | 6000 | 7000 | 8000 | 9000 | 10000 |
| --- | --- | --- | --- | --- | --- | --- | --- | --- | --- | --- |
| NaRnEA | 5641 | 5641 | 5641 | 5641 | 5641 | 5641 | 5641 | 5641 | 5641 | 5641 |
| GEP | 4291 | 4520 | 4674 | 4755 | 4831 | 4893 | 4942 | 4982 | 5038 | 5064 |

